# Quantitative Genetics of Microbiome Mediated Traits

**DOI:** 10.1101/2024.12.16.628599

**Authors:** Bob Week, Peter L. Ralph, Hannah F. Tavalire, William A. Cresko, Brendan J. M. Bohannan

## Abstract

Multicellular organisms host a rich assemblage of associated microorganisms, collectively known as their “microbiomes”. Microbiomes have the capacity to influence their hosts’ fitnesses, but the conditions under which such influences contribute to evolution are not clear. This is due in part to a lack of a comprehensive theoretical framework for describing the combined effects of host and associated microbes on phenotypic variation. Here we address this gap by extending the foundations of quantitative genetic theory to include host-associated microbes, as well as alleles of hosts, as factors that explain quantitative host trait variation. We introduce a way to partition host-associated microbiomes into components relevant for predicting a microbiome-mediated response to selection. We then apply our general framework to a simulation model of microbiome inheritance to illustrate principles for predicting host trait dynamics, and to generalize classical narrow and broad sense heritabilities to account for microbial effects. We demonstrate that microbiome-mediated responses to host-level selection can arise from various transmission modes, not solely vertical, but that the contribution of non-vertical modes can depend strongly on host life history. Our work lays a foundation for integrating microbiome-mediated host variation and adaptation into our understanding of natural variation.

## 1 INTRODUCTION

### 1.1 Overview

Nearly every lineage of multicellular organisms has a complex assemblage of associated microorganisms. This “microbiome” can contribute important physiological and developmental functions to their hosts (Spor et al., 2011; Burns et al., 2017; Goodrich et al., 2016; McFall-Ngai, 2007). Indeed, evidence that microbiomes play a vital role in shaping host fitness across a broad range of taxa is rapidly emerging (Argaw-Denboba et al., 2024; Houwenhuyse et al., 2021; Petersen et al., 2023; Gould et al., 2018; O’Brien et al., 2023; Rosshart et al., 2017). Although recent work has greatly expanded our understanding of the dynamics and function of host-associated microbiomes (Bruijning et al., 2021; Theis, 2018; Rosenberg and Zilber-Rosenberg, 2018; Roughgarden, 2023; Sandoval-Motta et al., 2017a), it is still unclear the extent to which microbiomes contribute to host evolution, and how such contributions depend on the mode of microbial transmission (Wendling and Wegner, 2015; Zilber-Rosenberg and Rosenberg, 2008; Russell, 2019). This knowledge gap is due in part to the lack of a comprehensive theoretical framework for modeling how host-microbiome system-level traits emerge from the interactions of host genes, microbiome and environment (Theis, 2018; Martiny et al., 2015; van Vliet and Doebeli, 2019; Roughgarden et al., 2017; Zilber-Rosenberg and Rosenberg, 2008).

Recently there have been calls to develop such a theoretical framework by incorporating microbiomes into quantitative genetics (Mueller and Linksvayer, 2022; Henry et al., 2021; Awany and Chimusa, 2020; Benson et al., 2010; Wang et al., 2018), a rich mathematical theory that models the inheritance and evolution of complex traits (Lynch and Walsh, 1998; Mackay et al., 2009; Rice, 2004; Walsh and Lynch, 2018). While microbiomes have sometimes been treated as traits influenced by host genetics (Camarinha-Silva et al., 2017; Knights et al., 2014; Opstal and Bordenstein, 2015), they have only recently been considered as inherited contributors to variation of emergent host–microbiome phenotypes (Org et al., 2015; Sandoval-Motta et al., 2017a). This shift has encouraged the use of microbiome-wide association studies (MWAS) by analogy to genome-wide association studies (GWAS) (Camarinha-Silva et al., 2017; Difford et al., 2018; Qin et al., 2012; Pérez-Enciso et al., 2021). However, caution is warranted: microbiomes are dynamic ecological assemblages obeying different rules of inheritance than genomes, and so uncritical application of methods designed for GWAS may produce misleading results. Microbiomes are especially challenging to incorporate into quantitative genetics because their inheritance is not Mendelian, and microbes need not be vertically transmitted (Bruijning et al., 2021; Uller and Helanterä, 2013; Roughgarden, 2023; van Vliet and Doebeli, 2019). Taken together, a careful extension of quantitative genetics is needed that accounts for the fundamental differences between host genetic and microbial sources of variation. Such an extension will allow theoretical investigations into how different modes of microbial transmission influence the evolution of host and microbial joint phenotypes, and will help clarify the degree to which these processes can affect long-term evolutionary trajectories.

Another body of theoretical work that considers host-associated microbiomes is the broad framework of inclusive inheritance (Day and Bonduriansky, 2011; Bonduriansky and Day, 2009; Uller and Helanterä, 2013; Helanterä and Uller, 2010; Danchin et al., 2011; David and Ricard, 2019). In this view, direct parent–offspring microbiome transmission can be treated as a maternal effect (Kirkpatrick and Lande, 1989), social transmission as an indirect genetic effect (Moore et al., 1997; Bijma et al., 2007), and environmental acquisition as a form of ecological inheritance (Odling-Smee et al., 2013). These perspectives have been explored in recent work (Week et al., 2024), which emphasizes that while each captures important aspects of microbiome-mediated evolution, none provides a complete account of microbiome inheritance dynamics. The unified framework introduced by Day and Bonduriansky (2011) offers a powerful theoretical foundation for integrating multiple inheritance modes into a single process-based model. However, this approach relies on explicit assumptions about transmission mechanisms, fidelity, and mutability parameters that remain poorly characterized or system-specific in many microbiome contexts, limiting its usefulness as a general foundation for studying host–microbiome evolution. This underscores the need for statistical frameworks that characterize patterns of microbiome inheritance, quantify their contribution to host trait variation, and predict microbiome-mediated responses to selection without presupposing detailed transmission rules.

Previous theoretical work focusing on host–microbiome evolution has yielded valuable insights, yet important gaps remain in our understanding of how such interactions contribute to complex host phenotypes and adaptation. For instance, Henry et al. (2021) emphasize that host trait variance can, in principle, be partitioned into components attributable to host genetics and the host-associated microbiome. However, this partitioning has not been formally defined, nor are quantitative predictions provided. At the other end of the spectrum, Zeng et al. (2017) introduce a mechanistic metacommunity model that integrates microbiome dynamics with host evolution. While this work offers valuable insights into the ecological foundations of microbiome inheritance, it focuses on host fitness rather than arbitrary traits, assumes all host variance arises from the microbiome, and considers only direct parent-offspring and environmental transmission. Roughgarden (2023) has proposed a general conceptual framework for microbiome inheritance, introducing the notions of lineal and collective inheritance. Lineal inheritance maintains correlations between parents and offspring at the individual level (e.g., via direct or intimate neighborhood transmission), whereas collective inheritance maintains correlations between generations at the population level (e.g., via social or indirect environmental transmission). However, this framework adopts a population genetic perspective and does not provide a means to partition host trait variation, and linking patterns of host-microbe ancestral overlap to the microbiome-mediated component of host trait variance is essential for predicting microbiome-mediated responses to selection on host populations. Taken together, these limitations point to the need for a framework that quantifies microbiome-mediated trait variance and links it to host-microbe ancestry relationships that underlie patterns of microbial inheritance.

In this study, we aim to complement prior work by (i) formally partitioning host trait variance into components attributable to host genetics and the microbiome, extending the classical approach of Fisher (1918); (ii) introducing retrospective, microbe-centered criteria to classify microbes based on concordance of their ancestry with host ancestry, complementing the inheritance concepts of Roughgarden (2023); and (iii) combining these tools to further decompose the microbiome component of host trait variance and to derive novel quantitative predictions for the response of microbiome-mediated traits to host-level selection. By categorizing microbes based on patterns of shared ancestry, our framework is independent of specific mechanistic assumptions on the modes of microbiome transmission. Given that patterns of microbe-host ancestral concordance are potentially measurable, this provides a unique approach to predict microbiome-mediated host adaptation without detailed knowledge of the underlying transmission mechanism. Additionally, our framework is flexible with respect to input data, accommodating coarse indices of microbial taxon presence/absence or relative abundances to fine-scale microbial SNPs. Our variance decomposition naturally generalizes additive genetic variance, separating contributions from host genetics, distinct microbiome components, and their covariances. Using simulations, we explore the conditions under which different components of the microbiome are expected to mediate a response to selection, and propose extensions of narrow- and broad-sense heritabilities that quantify microbiome-mediated contributions to host-level evolutionary change.

The remainder of the paper proceeds as follows. We begin by reviewing foundational principles from quantitative genetics, which ground our theoretical development. We then introduce our framework for partitioning host trait variance and classifying microbial ancestry, followed by simulations that explore the conditions under which different components of the microbiome are expected to mediate a response to selection. Finally, we extend classical heritability concepts by defining microbially informed analogues of narrow- and broad-sense heritability, and introduce associated transmissibility indices that quantify the evolutionary impact of distinct microbial components.

### 1.2 Quantitative genetic foundational principles

Quantitative genetics developed in the late 19th century through the development of statistical models to describe how continuous phenotypic variation depended on heritable and environmental components and to predict the phenotypic response to selection (Walsh and Lynch, 2018). Even after the rediscovery of Mendel’s laws in 1904, debate continued as to how continuous variation could emerge from discrete genes (Provine, 2001). R.A. Fisher settled the issue in a foundational paper (Fisher, 1918) showing that distributions of continuous phenotypes arise when alleles at multiple loci are considered. Following this, Fisher then introduced the concepts of *average excess* and *average effect* of allele substitutions in order to associate an additive quantity with each allele that best predicts the response to selection (Bürger, 2000; Fisher, 1941). This general approach follows the widely used theory of multivariate linear models, a perspective we use in extending the theory to account for microbial factors. Because these concepts are at the core of theoretical quantitative genetics, we summarize the basic principles here.

For a given host genotype *g*, write *z*_*g*_ as the host trait associated with *g* averaged over all other trait mediating factors (including environmental factors and host microbiome composition). The quantity *z*_*g*_ is referred to as a *genotypic value*. Then, writing *p*_*g*_ for the frequency at which host genotype *g* occurs in the population, we have that 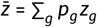 is the average of *z*_*g*_ over all host genotypes (equivalent to the population mean trait since all other factors have already been averaged over), and the quantity 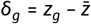 is the *average excess* associated with host genotype *g*. Fisher decomposed the average excess into an additive effect *α*_*g*_ (defined to be the sum of additive effects associated with each allele in host genotype *g*) and a residual deviation *ρ*_*g*_ such that *δ*_*g*_ = *α*_*g*_ + *ρ*_*g*_.

The definition of genotypic values implies the total host phenotypic variation *P* can be decomposed as *P* = *G* + *E*, where 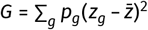 is the variance of genotypic values across the population (called the *genetic variance*). Although *E* is often referred to as the *environmental variance*, it is more precisely defined as *E* = *P* − *G*, the remaining variance left unexplained by genotypic values. Furthermore, defining the *additive genetic variance* 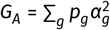 and residual variance 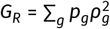, and defining the additive effects *α*_*g*_ to be chosen to minimize *G*_*R*_ (subject to the additivity constraint); with this choice the genetic variance decomposes as *G* = *G*_*A*_ + *G*_*R*_ (Bürger, 2000). Although (Fisher, 1918) assumed Hardy-Weinberg equilibrium, which allowed derivation of more specific results, the overall approach only depends on the identification of the genotypic values *z*_*g*_.

This definition of *G*_*A*_ turns out to be useful for accurately predicting the response of a trait to selection. For instance, *G*_*A*_ is more useful than *G* in predicting the response to selection because parents transmit alleles, not entire genotypes. In particular, the classical *breeders equation* takes the form 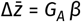, where 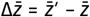 is the difference between offspring mean trait 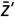 and parent mean trait 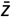, and *β* = Cov(*w, z*)/*P* is the correlation coefficient of fitness (*w*) with phenotype (Lande, 1976).

These definitions are also used to define *heritability* of a trait as a measure of parent-offspring resemblance. In particular, the quantities *h*^2^ = *G*_*A*_/*P* and *H*^2^ = *G*/*P* are referred to as the *narrow* and *broad* sense heritabilities, respectively (reviewed by Visscher et al., 2008). With this notation, we note that the breeders equation was initially introduced by Lush (1937) in the form *R* = *h*^2^*S*, where 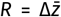 is the response to selection and *S* is the selection differential (see Walsh and Lynch, 2018). Key notation used throughout our paper is summarized in Table 1.

**TABLE 1.**
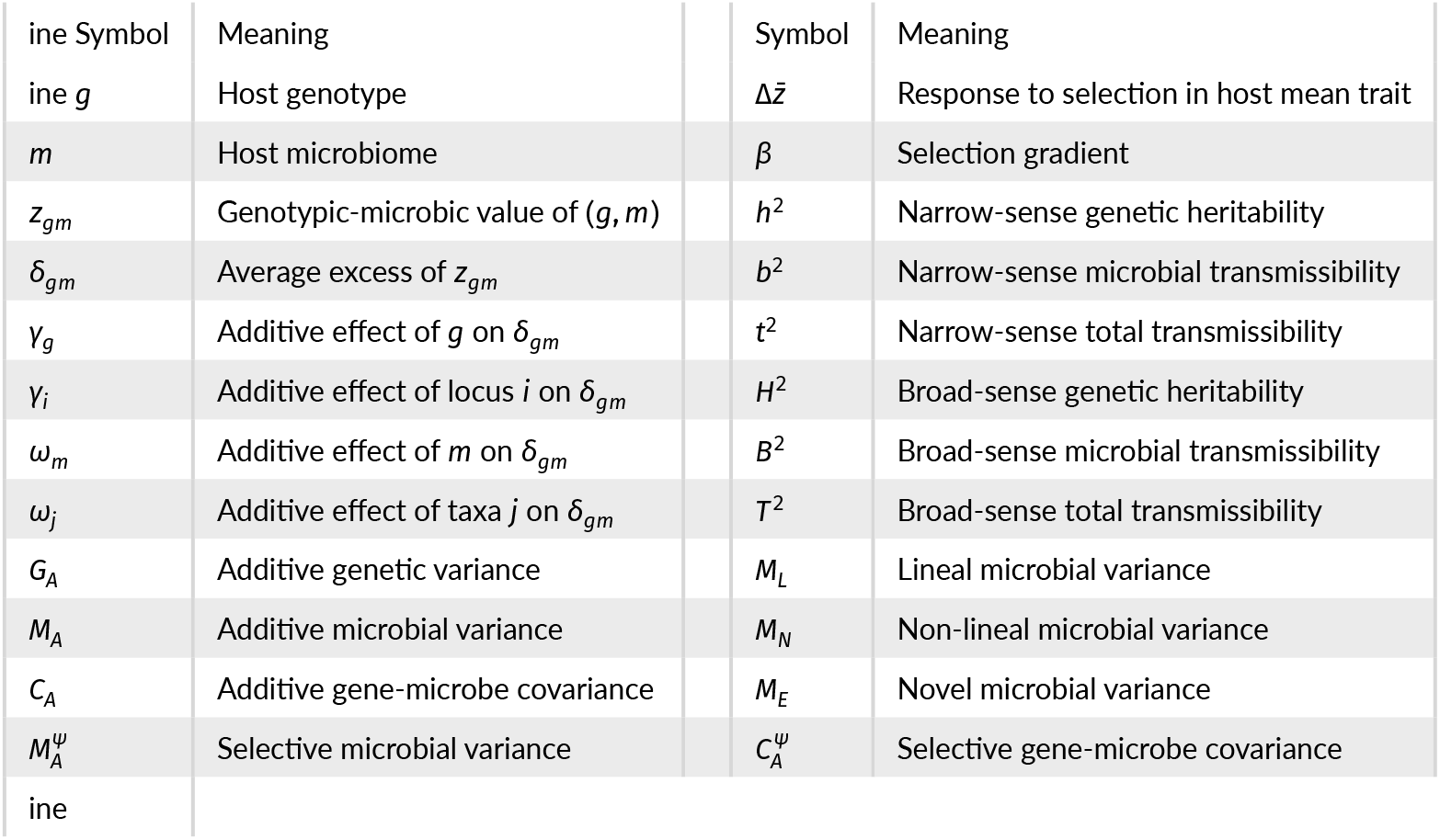
Summary of key notation.

In what follows we build on these foundations to incorporate the effects of host-associated microbiomes. By generalizing the above approach, we naturally arrive at a generalization of additive genetic variance, the breeders equation, and narrow-sense and broad-sense heritabilities for microbiome mediated traits. Before we introduce our framework, we discuss some initial assumptions for incorporating microbiomes in the following section.

### 1.3 Perspectives on incorporating microbes into host evolution

By building on quantitative genetics theory we are admittedly using a very host-centric framework that leads to several constraining assumptions. In particular, we focus on a single trait that is an emergent property of the host-microbe system, such as the ability of the host to acquire energy from food that first must be processed by components of the host gut microbiome. For example, termites and their associated gut microbiomes together can process high amounts of cellulose to acquire energy (Maurice and Erdei, 2018; Ali et al., 2019; Arora et al., 2022).

We also consider contributions from both host genes and individual microbes to this trait as potentially heritable units, meaning that they can be transmitted from a donor to a recipient, and influence the value of the emergent phenotype. Traditional quantitative genetics defines the donor and recipients solely as parents and offspring through the transmission of alleles. In sexually reproducing diploid populations, the process of gametogenesis and syngamy leads to Mendelian segregation.

In contrast, microbes can be transmitted among hosts in a variety of ways. In particular, a microbial parent (donor) and microbial offspring (recipient) may not be genetically related. Additionally, unlike the transmission of alleles via sexual reproduction, microbial recipients can have more than two donors, depending on the microbial taxa considered and mode of transmission. In some cases the donor and receiver of transmitted host genetic and microbial units will be the same, which has been described as *lineal transmission* (Roughgarden, 2023) and *co-propagation* (Mueller and Linksvayer, 2022). In these cases, narrow sense heritability of microbiome-mediated traits may be confounded by microbiome inheritance, possibly inflating estimates of heritability based on pedigree compared to estimates based on SNPs (Sandoval-Motta et al., 2017b).

We must also take into account that microbiome composition is governed by community assembly and not Mendelian processes, and microbial abundances can vary dynamically throughout the life of the host. These have been termed the *fidelity of transmission* and *persistence fidelity* of microbes (Mueller and Linksvayer, 2022), with the idea that microbe persistence is often less than that of an allele. Based on this consideration, one possible path forward is to study models of metacommunity dynamics as the basis of microbiome inheritance and fidelity from the lens of an extended framework of quantitative genetics (Leibold et al., 2004; Koskella et al., 2017; Zeng et al., 2017; Miller et al., 2018). Here we focus on beginning this extension of quantitative genetics. Analysis of metacommunity models of microbiome inheritance using this framework is outside the scope of this paper, but it is likely microbiome dynamics will alter indices of microbiome-mediated host trait inheritance.

Several authors have recently attempted to extend heritability to include microbial contributions, and have proposed novel terminology such as *microbiability* and *holobiontability*. However, we follow the convention established by Rothschild et al. (2018) and clarified by Mueller and Linksvayer (2022), who use *H*^2^ and *h*^2^ for broad- and narrow-sense *genetic heritability*, and *B*^2^ and *b*^2^ for what they term *microbiome heritability*. To avoid conflating genetic and microbial sources of variation, we refer to the latter as *microbial “transmissibility*.*”* As Mueller and Linksvayer (2022) make clear, *B*^2^ and *b*^2^ refers to microbial contributions to the emergent host phenotype, not to be confused with the genetic heritability of microbiomes as host traits in and of themselves (Morris and Bohannan, 2024). Here we propose a more precise usage of this terminology to refer specifically to the proportion of host trait variance mediated by the component of host microbiome that is both transmitted across host generations and in some cases mediates a response to host-level selection (see section 6 below).

## 2 MATERIALS AND METHODS

### 2.1 Analysis of variance of microbiome mediated traits

The partitioning of variance reviewed in section 1.2 is not restricted to genetic material, but can be applied to any collection of trait mediating factors (e.g., soil salinity, local rainfall, temperature, oxygen concentration, pH, etc.). In particular, for a microbiome mediated host trait, relevant factors include the host genotype *g* and host microbiome *m*. The overall approach is sufficiently general to accommodate any summary of host genotype and host microbiome, such as presence/absence of microbial taxa and microbial SNPs obtained from 16s data or shotgun metagenomes. However, for the sake of clarity, we describe our framework focusing on host allele counts and relative abundances of microbe taxa.

We consider diploid host populations with *L* biallelic loci. We denote by *g*_*i*_ ∈ {0, 1, 2} the number of alleles at locus *i* ∈ {1, …, *L*} (the choice of allele counted is arbitrary). Turning to the host microbiome, we assume microbiomes may be summarized by the relative abundances of *S* microbial taxa (relative to the total microbiome abundance) with *m*_*j*_ being the relative abundance of microbe taxon *j* ∈ {1, …, *S*}. Vectors of allele counts and relative microbial abundances are respectively written as *g* = (*g*_1_, …, *g*_*L*_)^†^ and *m* = (*m*_1_, …, *m*_*S*_)^†^, where ^†^ denotes matrix transposition.

We write *z*_*gm*_ as the expected trait value for hosts carrying the genotype-microbiome pair (*g, m*) averaged across all other trait mediating factors, and refer to *z*_*gm*_ as the *genotypic-microbic value* of (*g, m*). Additionally, we write 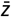 as the mean *genotypic-microbic value* across all hosts in the population so that 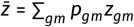, where *p*_*gm*_ is the frequency of hosts carrying the genotype-microbiome pair (*g, m*) and the sum is taken over all possible combinations of (*g, m*). By definition,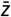 is also the mean trait of the host population.

The *average excess* of the pair (*g, m*) is then 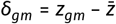, and we consider the decomposition *δ*_*gm*_ = *α*_*gm*_ + *ρ*_*gm*_, where *α*_*gm*_ is an additive component that decomposes into additive genetic and additive microbial effects of the pair (*g, m*) on *δ*_*gm*_, and *ρ*_*gm*_ is the residual deviation. In particular, following Fisher’s general approach, we assume *α*_*gm*_ = *γ*_*g*_ + *ω*_*m*_ where *γ*_*g*_ is the additive genetic effect and *ω*_*m*_ is the additive microbial effect on the average excess *δ*_*gm*_.

Finally, using our assumptions for summarizing host genotype and host microbiome, the definition of additive effects imply that they can be written as

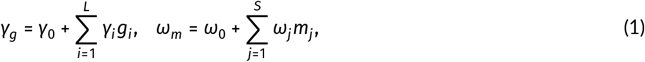

where *γ*_*i*_ is the per count additive allelic effect of locus *i* ∈ {1, …, *L*} on *δ*_*gm*_ and *ω*_*j*_ is the additive effect of the relative abundance of taxon *j* ∈ {1, …, *S*} on *δ*_*gm*_. To be clear, *γ*_*g*_ is distinguished from *γ*_*i*_ because *g* is a vector and *i* is an integer. A similar distinction holds for *ω*_*m*_ and *ω*_*i*_. Lastly, the sum *α*_0_ = *γ*_0_ + *ω*_0_ acts as an intercept for the linear model, and the components *γ*_0_ and *ω*_0_ will be determined by our decomposition of *P*_*A*_ below.

Following the general scheme outlined in section 1.2, we can write the additive component of host trait variation explained by host genetic and host microbiome factors as 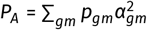. The definition of *P*_*A*_ is formalized by solving the least squares problem of minimizing 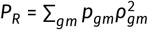 with respect to *α*_0_, *γ*_1_, …, *γ*_*L*_, *ω*_1_, …, *ω*_*S*_, where *ρ*_*gm*_ = *δ*_*gm*_ −*α*_*gm*_. In supplement section 1 we show that setting Cov(*z*_*gm*_, *g*) to the vector with *i*th entry Cov(*z*_*gm*_, *g*_*i*_), Cov(*z*_*gm*_, *m*) to the vector with *j*th entry Cov(*z*_*gm*_, *m*_*j*_), Γ to the matrix with *ij*th entry Cov(*g*_*i*_, *g*_*j*_), Ω to the matrix with *ij*th entry Cov(*m*_*i*_, *m*_*j*_), and Ξ to the matrix with *ij*th entry Cov(*g*_*i*_, *m*_*j*_), the least-squares problem reduces to solving the linear system

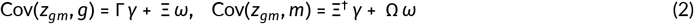

for *γ* = (*γ*_1_, …, *γ*_*L*_)^†^ and *ω* = (*ω*_1_, …, *ω*_*S*_)^†^, where ^†^ denotes matrix transposition. The solution is then given by

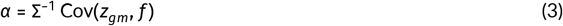

with *α* = (*γ*_1_, …, *γ*_*L*_, *ω*_1_, …, *ω*_*S*_)^†^, *f* = (*g*_1_, …, *g*_*L*_, *m*_1_, …, *m*_*S*_)^†^, 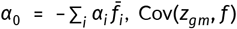 the vector with *i*th entry Cov(*z*_*gm*_, *f*_*i*_), and Σ the block matrix

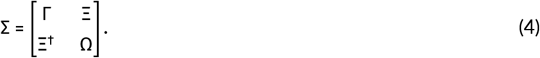

Similar to the case for additive genetic effects reviewed in section 1.2, this solution is unique so long as Σ has an inverse. This means that if the allele counts at any pair of loci, or if the relative abundances of any pair of microbes, or if an abundance of a microbe and an allele count of a locus perfectly correlate, the solution is no longer unique.

Since we have centered the trait (the average of *δ*_*g*_ is zero), it turns out that with *α*_0_, *γ*, and *ω* as above, that 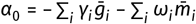, where 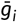 is the average number of alleles at locus *i* (which is twice the allele frequency due to host genotypes being diploid) and 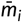 is the average relative abundance of microbe taxon *i*. Then, it is convenient to set 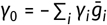 and 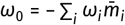. With these choices for *γ*_0_ and *ω*_0_, the additive components of host trait variance can be expressed in terms of allele counts and relative microbe abundances as:

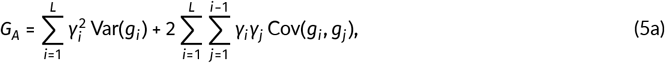

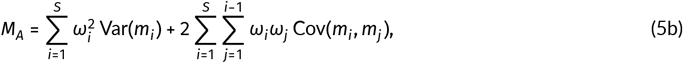

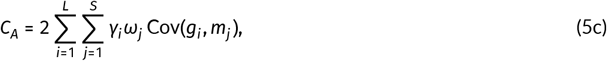

where we use the convention that an empty sum 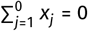 is zero.

More pragmatically, these expressions can be obtained from a least-squares fit of a multivariate linear model that predicts a sample of traits *ζ* = (*z*_1_, …, *z*_*n*_)^†^ from associated samples of host gene sequences and microbiome compositions (collected into the data matrix *X* which, for its *i*th row, has allele counts at *L* loci and relative microbe abundances across *S* taxa associated with the *i*th sample) such that *ζ* = *X α* + *ε*, with *ε* = (*ε*_1_, …, *ε*_*n*_)^†^ being the error terms for each sample. Various algorithms exist for efficiently finding a least-squares solution for the additive effects *α*, such as pivoted QR decomposition (Strang et al., 2019).

This overall approach holds for genotypic-microbic values occurring as arbitrary functions of genotype-microbiome pairs; *z*_*gm*_ = *φ*(*g, m*) for any function *φ*. That is, this approach accounts for any kind of interaction among genes, among microbes, and between genes and microbes on the expression of host traits. To illustrate, we apply this approach to a model of genotypic-microbic values that includes interactions between genes and microbes in the following section.

### 2.2 A model of gene-microbe interactions

The approach to analysis of variance above establishes a statistical model. We can then apply this model either to empirical or simulated data. As a demonstration, we examine what the statistical model tells us when applied to a mechanistic model involving pairwise interactions between host loci and microbial taxa. Considering *z*_*gm*_ = *φ*(*g, m*) as a function of the genotype-microbiome pair (*g, m*), we analyze the model

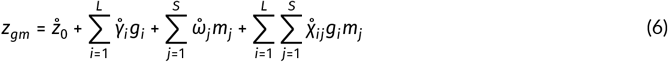

where 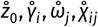 are the mechanistic model parameters, and 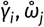 are not necessarily equivalent to the additive effects *γ*_*i*_, *ω*_*j*_ as we now show.

Using the approach outlined in the previous section, the additive genetic and microbial effects from model (6) depends on Cov(*z*_*gm*_, *g*_*k*_) and Cov(*z*_*gm*_, *m*_*k*_), which are given by

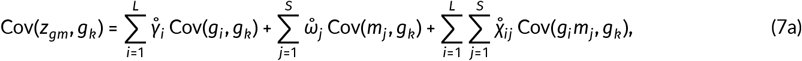

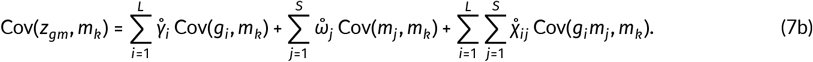

Writing

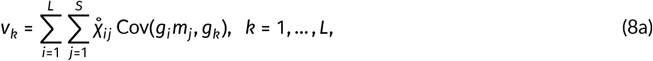

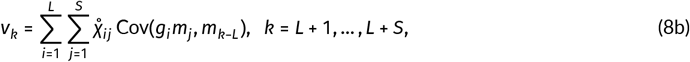

the additive genetic and microbial effects are given by 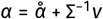, where *α* = (*γ*_1_, …, *γ*_*L*_, *ω*_1_, …, *ω*_*S*_) are the additive genetic and microbial effects, Σ is defined at the end of the previous section, and 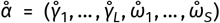 are mechanistic model parameters.

In the absence of gene-microbe interactions, so that 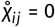 for all *i* = 1, …, *L* and *j* = 1, …, *S*, we have *v* = 0 and the additive effects formally defined above become equivalent to the additive effects of the model; 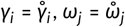. However, if 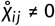 for any host locus *i* and any microbial taxon *j*, the formal additive effects and model additive effects are in general no longer equivalent 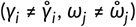 because the formal additive effects will also include terms due to gene-microbe interactions.

To unpack this more, consider the example where the effect of a microbe only occurs in the presence of an allele at a haploid biallelic locus such that neither the allele nor the microbe have an effect on the host trait in the absence of the other. In this case, the above model simplifies to 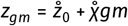, where *g* = 0, 1 determines the presence of the allele, *m* = 0, 1 determines the presence of the microbe, and 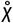 quantifies the effect of their interaction. Writing *p*_*gm*_ as the frequency of hosts carrying the pair (*g* = 1, *m* = 1), *p*_*g*_ the frequency of hosts with the allele (so the marginal frequency of *g* = 1), and *p*_*m*_ the frequency of hosts with the microbe (so the marginal frequency of *m* = 1), then the additive effect of the allele is *γ ∝* (*p*_*m*_ − *p*_*gm*_)(1 − *p*_*m*_), and the additive effect of the microbe is *ω ∝* (*p*_*g*_ − *p*_*gm*_)(1 − *p*_*g*_), where *∝* means *proportional to* and the constant of proportionality for both quantities is 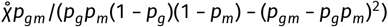. Hence, even in the absence of any underlying additive effects in the mechanistic model, the formally defined additive effects of the statistical model are non-zero. Additionally, each of these effects depends on the frequency of the other factor.

The observation that non-additive mechanistic effects can contribute to additive components of trait variance directly extends the recognition that epistasis contributes to additive genetic variance (Cheverud and Routman, 1996; Barton and Turelli, 2004). In supplement section 2 we provide expressions for additive genetic and additive microbial effects given a model that includes all possible pairwise interactions between genes and microbes explained by products of allele counts and microbial abundances.

### 2.3 Microbiome mediated response to host-level selection

Because the transmission of microbes is not necessarily direct from parent to offspring, nor does it occur with high fidelity relative to genetic transmission (e.g., microbiome composition exhibits significant variation between related individuals, see Tierney et al., 2019; Tavalire et al., 2021), we cannot transfer many of the important simplifying assumptions of classical quantitative genetics regarding inheritance to the study of microbiome mediated traits. These simplifying assumptions have also been central to the power of the classical quantitative genetic approach, such as the use of pedigree analysis in the animal model (Wilson et al., 2009). Additionally, the degree to which additive host trait variation explained by microbes mediates an inter-generational response to selection on host individuals is not obvious. To overcome these issues, and identify components of host microbiomes that contribute host population-level selection responses, we propose a classification scheme to account for different patterns of ancestral concordance between microbes and the hosts that carry them. We then apply a set of simplifying assumptions to clarify our initial analysis, and consider consequences for a microbially mediated heritable response to selection on host individuals. This analysis motivates definitions of microbial heritabilities and host trait transmissibilities that we introduce below in section 3.2.

#### 2.3.1 Classifying microbes by concordance with host ancestry

As a first step in addressing the challenge described above, we classify host-associated microbes according to the ancestral relationship between their lineages and those of the hosts they inhabit. Specifically, we consider the degree to which microbial lineages are associated with host lineages. A lineage is a chain of (genetic) parent-offspring relationships, and so the distinction regards the degree to which transmission of microbes matches transmission of genetic material. These definitions make no assumptions about transmission mode (e.g., social, environmental, parent-offspring). This leads to the following partition:

1. **Lineal ancestry:** Microbes are classified as *lineal* if their lineages have historically remained associated with the same host lineage across multiple generations (Figure 1, left panel). In other words, if the ancestor of a given microbe within an individual host at some point back in time can be found in a host who is a direct (genealogical) ancestor of that individual. These microbes are most likely to have co-diverged, co-adapted, or co-transmitted with their host lineage and are therefore most likely to contribute to selection responses at the host population level.
2. **Non-lineal ancestry:** Microbes are classified as *non-lineal* if their lineages have been associated with host lineages but do not consistently follow host lineages (Figure 1, center panel). For instance, these microbes may frequently disperse horizontally between related hosts such as siblings, or between less related conspecific hosts, or may move back and forth between hosts and the environment. Their ancestry is partially concordant with host ancestry, and their contribution to host adaptation may vary depending on transmission routes and patterns of host association.
3. **Novel ancestry:** Microbes are classified as *novel* if their ancestral lineages have not spent any time with host ancestors (Figure 1, right panel). These microbes originate from sources external to the host population (e.g., biotic or abiotic environmental reservoirs) and represent recently acquired microbial lineages with no history of host association. By definition, novel microbes are not inherited through host lineages and are unlikely to have experienced selection aligned with host reproductive success. All novel microbes are by definition environmentally acquired, but not all environmentally acquired microbes are novel.

These definitions, formulated at the scale of microbial individuals, are illustrated in Figure 1. The goal of this scheme is to partition host microbiomes into components that are (1) unlikely to contribute to host population–level selection responses (novel microbes), (2) most likely to do so (lineal microbes), or (3) of uncertain contribution (non-lineal microbes). For some host-microbiome systems, it may be necessary to add additional categories, for instance, between non-lineal and novel for microbes that are nearly novel, but with ancestral lineages that have spent significant time in the host population. This partitioning of host microbiomes is conceptually similar to the notions of lineal and collective inheritance introduced by Roughgarden (2023). Indeed, a lineal microbe in our framework would be considered lineally inherited in Roughgarden’s framework. The primary distinction is that Roughgarden’s concepts focus on inheritance processes, but our partitioning focuses on categorizing microbes by their tendency to be inherited (possibly lineally, collectively, or not at all). For instance, the class of non-lineal microbes spans a broad spectrum of associations with the host; many may be transient and therefore have limited capacity to contribute to collective inheritance.

**FIGURE 1.**
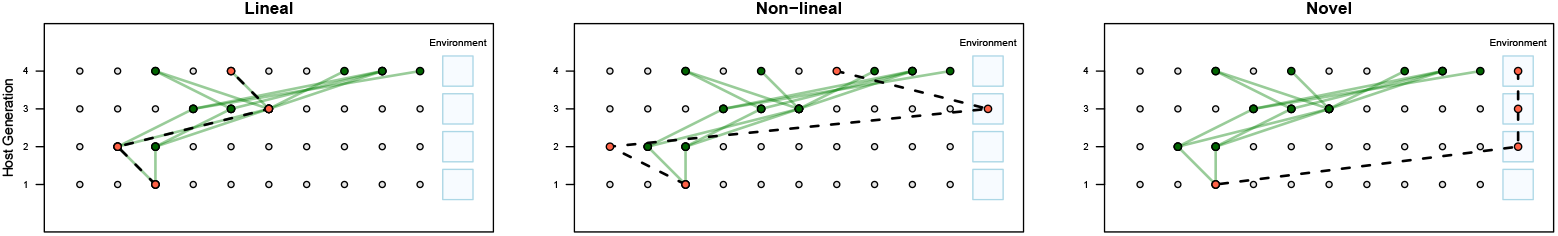
Illustration of our partitioning of host microbiomes by ancestral concordance. Each panel shows a different pattern of host-microbe ancestral concordance. Rows represent host generations (most recent at the bottom), with blue squares indicating the environment. Circular nodes are host individuals; green lines and circles trace the focal host’s ancestry, while dashed lines and tomato-colored circles trace the focal microbe’s lineage. **Left:** A lineal microbe whose ancestry closely tracks the host lineage: inheritance of microbes (dotted line) always follows genetic ancestry (green lines). **Middle:** A non-lineal microbe with ancestry mostly among hosts but not following genetic ancestry, and occasionally entering the environment. **Right:** A novel microbe whose ancestry lies entirely outside the host population.

To leverage this approach for understanding how microbiomes mediate responses to selection, we next quantify the degree to which signals of novel, lineal, and non-lineal ancestry are reflected in each microbial taxon within a host individual. These values are can be averaged across host individuals to obtain a summary measure of ancestral concordance for each microbial taxon. Assessing host-microbe ancestral concordance at the level of microbial taxa is especially useful in our framework, as it aligns with the scale at which we quantify microbial effects on host traits.

#### 2.3.2 Partitioning host trait variance by ancestral concordance

Although many microbial taxa likely exhibit several patterns of ancestral concordance, for this initial inquiry we consider a simplified scenario where each taxon consists of individuals that are either entirely novel, lineal, or non-lineal. This simplifies the further partitioning host trait variance according to the partitioning of microbiomes proposed above, and this simplicity allows us to highlight the consequences for host population-level selection responses. More realistic models that account for the complexity of microbial transmission could be studied in future work. In particular, assuming the *S* microbial taxa in the host microbiome can be subdivided into *S*_*L*_ lineal taxa, *S*_*N*_ non-lineal taxa, and *S*_*V*_ novel taxa (so that *S* = *S*_*L*_ + *S*_*N*_ + *S*_*V*_), we can rewrite the additive microbial effect on the average excess 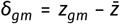 as *ω*_*m*_ = *𝓁*_*m*_ + *n*_*m*_ + *v*_*m*_ where

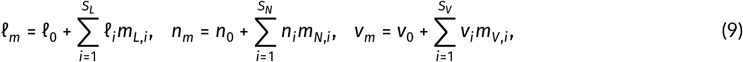

where the additional subscripts indicate the pattern of ancestral concordance (*L*ineal, *N*on-lineal, No*V*el) each group of taxa has with its host. Taking this partitioning a step further, we can write the additive microbial variance as

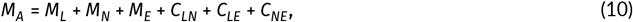

where 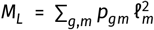 is the component of additive variance due to lineal microbes (which we refer to as the *additive lineal variance*), *M*_*N*_ = ∑_*g,m*_ *p*_*gm*_ *n*_*m*_ is the component due to non-lineal microbes (the *additive non-lineal variance*), *M*_*V*_ = ∑_*g,m*_ *p*_*gm*_ *v*_*m*_ is due to novel microbes (the *additive novel variance*), *C*_*LN*_ = 2 ∑_*g,m*_ *p*_*gm*_ *𝓁*_*m*_ *n*_*m*_ is due to non-random associations between lineal and non-lineal microbes (the *additive lineal-non-lineal covariance*), *CLV* = 2 ∑_*g,m*_ *p*_*gm*_ *𝓁*_*m*_ *v*_*m*_ is due to non-random associations between the abundances of lineal and novel microbes (the *additive lineal-novel covariance*), and *C*_*NV*_ = 2 ∑_*g,m*_ *p*_*gm*_ *n*_*m*_ *v*_*m*_ is due to non-random associations between the abundances of non-lineal and novel microbes (the *additive non-lineal-novel covariance*). For later use, we also introduce the *additive gene-lineal covariance C*_*L*_ = 2 ∑_*g,m*_ *p*_*gm*_ *γ*_*g*_ *𝓁*_*m*_ as the component of *additive gene-microbe covariance* due to lineal microbes, the *additive gene-non-lineal covariance C*_*N*_ = 2 ∑_*g,m*_ *p*_*gm*_ *γ*_*g*_ *n*_*m*_ as the component due to non-lineal microbes, and the *additive gene-novel covariance C*_*V*_ = 2 ∑_*g,m*_ *p*_*gm*_ *γ*_*g*_ *v*_*m*_ as the component due to novel microbes.

## 3 RESULTS

### 3.1 Interactions between host-level selection and ancestral concordance

By definition, novel microbes cannot contribute to a microbiome mediated response of the host trait to selection on hosts. In contrast, lineal microbes will likely make the most reliable contributions to a microbiome mediated response to host-level selection. The degree to which non-lineal microbes contribute to a microbiome mediated response depends on the the stage in host life-history that selection occurs relative to the stage in host life-history that non-lineal microbes are transmitted. For instance, if non-lineal microbes are first shed into the environment before the host parental population experiences host-level selection (so selection does not shape the set of non-lineal sources), then non-lineal microbes will not contribute to a microbiome mediated response. On the other hand, if non-lineal sources include only the microbiomes of host parents that, for example, survived an episode of viability-based selection, then non-lineal microbes may contribute significantly to a microbiome mediated response.

Based on the above reasoning, to obtain an accurate model for predicting a microbiome mediated response to selection, novel microbes must first be identified and culled from the set of factors considered, and their abundances should be averaged over when calculating *z*_*gm*_. Additionally, it must be determined whether or not non-lineal microbes are sourced from selected parents or from the broader population of unselected parents, because this is likely associated with whether or not they contribute to a response to selection. Having identified the microbial taxa that contribute to a response to selection, we write 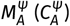 for the additive microbial variance (gene-microbe covariance) calculated using only the abundances of those taxa, and refer to this quantity as the *selective* additive microbial variance (gene-microbe covariance). Building on this logic, we anticipate three basic conclusions.

First, if we summarize the abundances of lineal microbes using presence/absence, and assume host offspring inherit these binary values for each taxon independently, then the inheritance of these microbes is structurally equivalent to the inheritance of genetic alleles at freely recombining loci. Hence, in this case, it is clear that lineal microbes can contribute to a response to selection. This clearly still holds when presence/absence is replaced with higher resolution summaries of microbial abundances such as approximated relative abundances. In particular, we expect the selective additive microbial variance 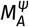 to always include the additive lineal variance *M*_*L*_, and the selective additive gene-microbe covariance 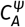 to always include the additive gene-lineal covariance *C*_*L*_ (defined at the end of section 2.3.2).

Second, if non-lineal microbes are sourced from the broader population of “unselected” parents (i.e., if the source of non-lineal microbes is independent of fitness), then we anticipate non-lineal microbes to not contribute to a response to selection. In this case the selective additive microbial variance would still simplify to the additive lineal variance,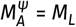, and the selective additive gene-microbe covariance would also still simplify as 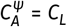.

However, as a third conclusion, if non-lineal microbes are sourced from selected parents, or more generally, values of the selected trait are correlated with probability of being a non-lineal source of microbes, we anticipate that their abundances will also contribute to a response to selection. In this case, selective additive microbial variation would become 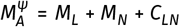 and selective additive gene-microbe covariance becomes 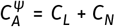. To test our expectations described above, we implemented a simulation model based on the general setup described at the beginning of section 2.1, which we now describe.

#### 3.1.1 Simulation description

Our simulation model assumes that host traits follow an additive model equivalent to that analyzed in section 2.2, except without any gene-microbe interactions (so 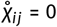 for each *i, j*). As a function of phenotype, host fitness is set to *W*(*z*) = exp(*sz*), which leads to directional selection for larger host traits when *s* > 0 (we set *s* = 10^−3^). We simulate 100 freely recombining host genetic loci. The abundances of lineal microbes in host offspring are Poisson distributed around the mid-parents of those taxa. For non-lineal microbes we consider two models: 1) each host offspring chooses a single host individual uniformly from the broader unselected parental population as its non-lineal source and 2) each host offspring chooses a single host individual from the selected parental population with probability proportional to its fitness. Once the non-lineal donor is chosen, the host offspring inherits non-lineal abundances that are Poisson distributed around the abundances of non-lineal taxa in the donor. Finally, novel microbe abundances are drawn independently and identically from a Poisson distribution for each taxa in each host in each host generation. We assume 100 microbial taxa in each microbiome component, with each taxa having an average abundance of 50 in each host parent. This model of microbial inheritance is illustrated in Figure 2.

**FIGURE 2.**
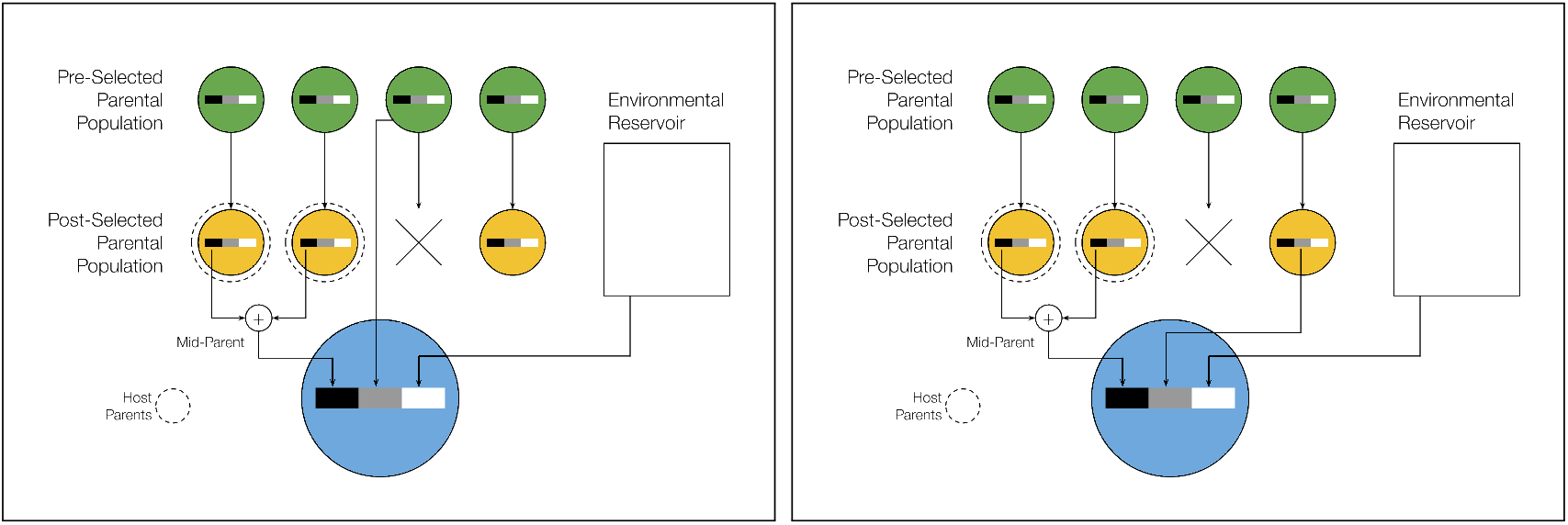
Graphical representation for the two models of microbiome inheritance used in our simulations. In the left panel, non-lineal microbes for a given offspring are acquired from a randomly chosen individual in the pre-selected parental population. In the right panel, non-lineal microbes are acquired from a randomly chosen individual in the post-selected population. Circles represent hosts and rectangles within hosts the lineal (black), non-lineal (gray), and novel (white) microbiome components. The green circles at the top of each panel represent the parental population before an episode of viability selection, and the mustard circles represent the parental population after an episode of viability selection (with vertical straight arrows indicating the trajectories of host individuals). The vertical rectangle on the right represents an environmental reservoir that novel microbes are sampled from. Hosts highlighted with dashed concentric circles are the parents of the focal offspring (the blue circle at the bottom). The plus symbol joining the lineal components of the parental microbiomes indicates taking an average. Arrows pointing towards offspring brackets represent microbiome transmission.

Results are obtained by first drawing normally distributed additive effects, and then observed and predicted responses to selection are averaged over randomly drawn genotype-microbiome pairs for host parents, repeated selection experiments, and repeated formation of offspring from selected parents. Figure 3 illustrates mean trait dynamics under our model when different combinations of factors mediate traits, and when non-lineal microbes are sourced from pre-selected or post-selected parents. In addition, time-series of correlations between host allele counts and microbe abundances are shown in Figure 4. Parameter values used for simulating data presented in Figures 3 and 4 are provided Table 1 of the supplement. Parameter values used for simulating data presented in Figure 5 are provided in Table 2 of the supplement. Code to reproduce these results (written in Julia) is provided at the GitHub repository https://github.com/bobweek/qgmmt.

**FIGURE 3.**
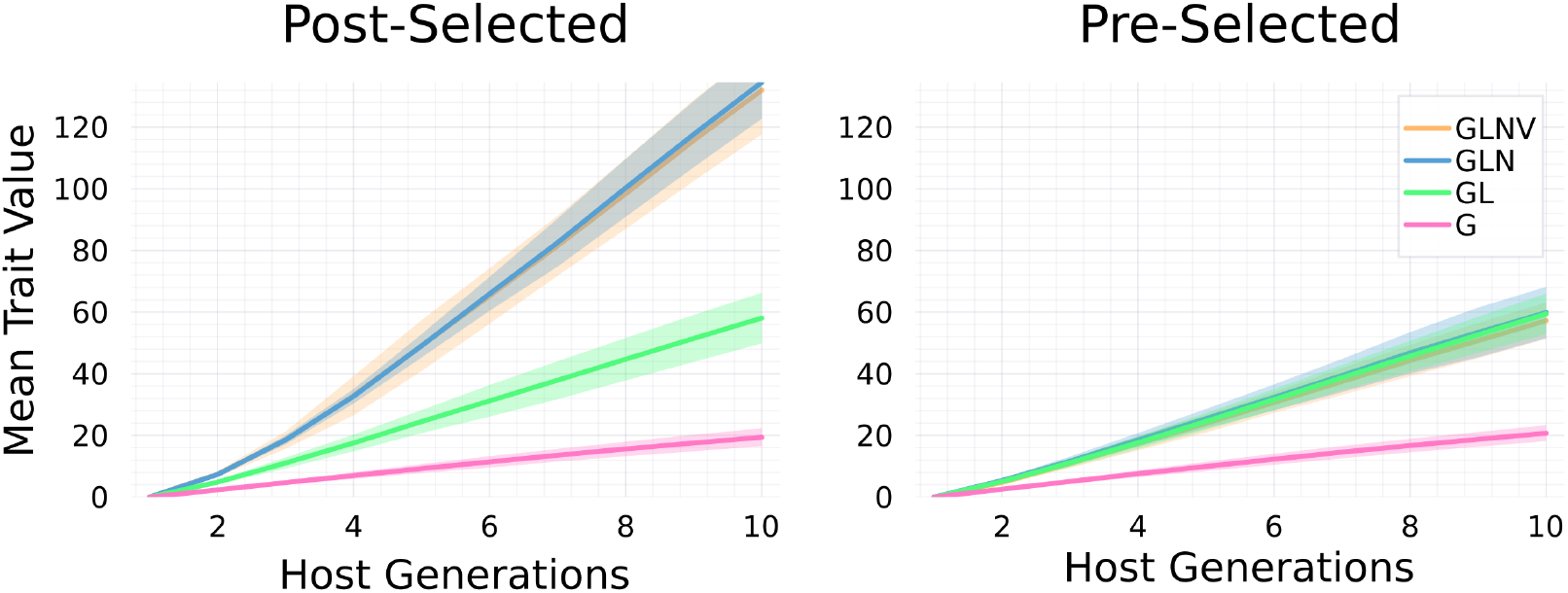
Time series plots of average simulated host trait dynamics. Different colors correspond to different combinations of factors. Pink corresponds to only host genes mediating a response to selection (G). Green corresponds to only host genes and lineal microbes (GL). Blue corresponds to host genes, lineal and non-lineal microbes (GLN). Mustard corresponds to host genes, lineal, non-lineal, and novel microbes (GLNV) (mostly hidden by blue lines/areas). For each combination of factors, simulations were repeated 20 times. Solid lines are averages across these repeated runs, and shaded regions correspond to standard deviations across repeated runs. The panels are divided by non-lineal microbes acquired from post-selected parents (left) and pre-selected parents (right).

**FIGURE 4.**
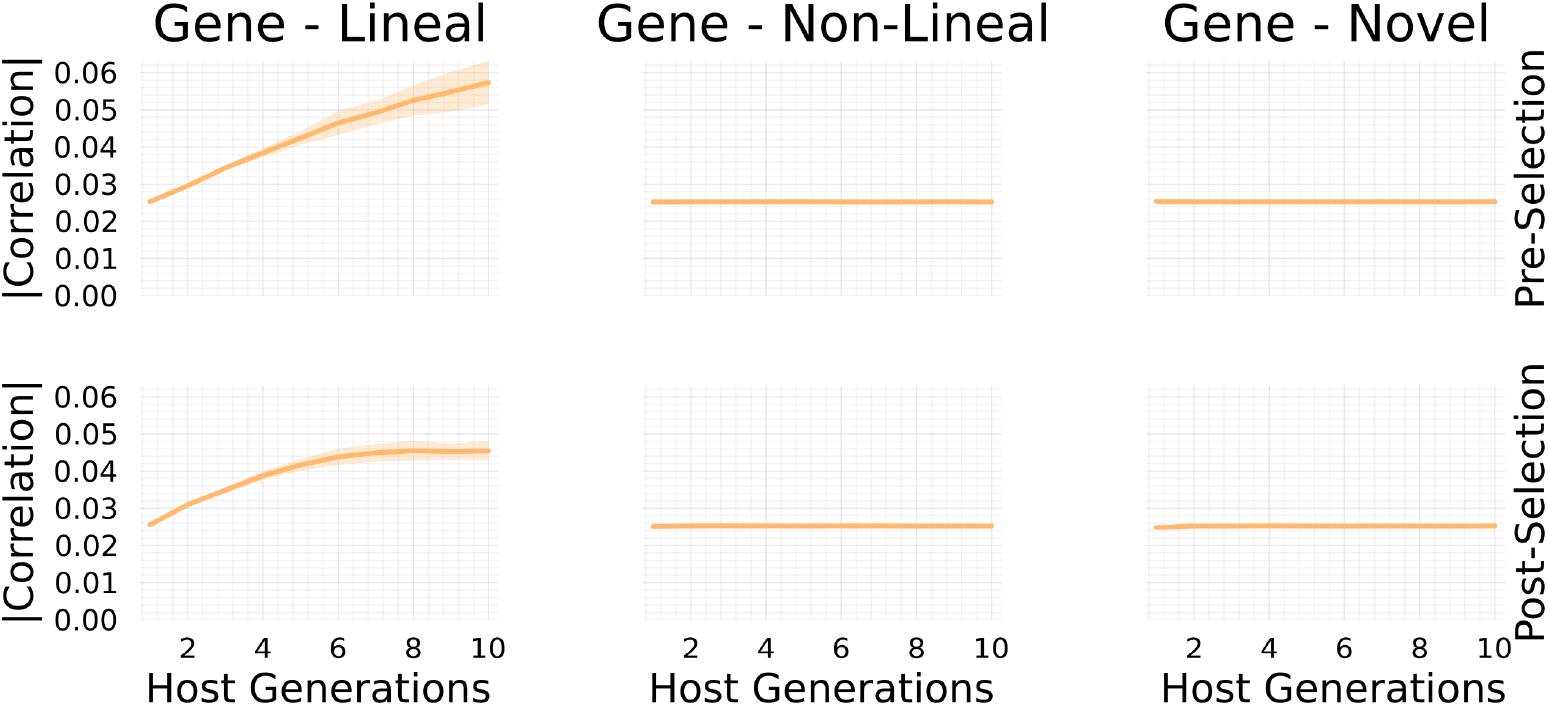
Time series plots of absolute values of correlations between host allele counts and microbe abundances. Correlations between allele counts and abundances of lineal microbes (left plots) increase faster when non-lineal microbes are sourced from the pre-selected host parental population in comparison to when non-lineal microbes are sourced from the post-selected host parental population. In contrast, correlations with abundances of non-lineal microbes and novel microbes remains at neutral levels (right plots).

**FIGURE 5.**
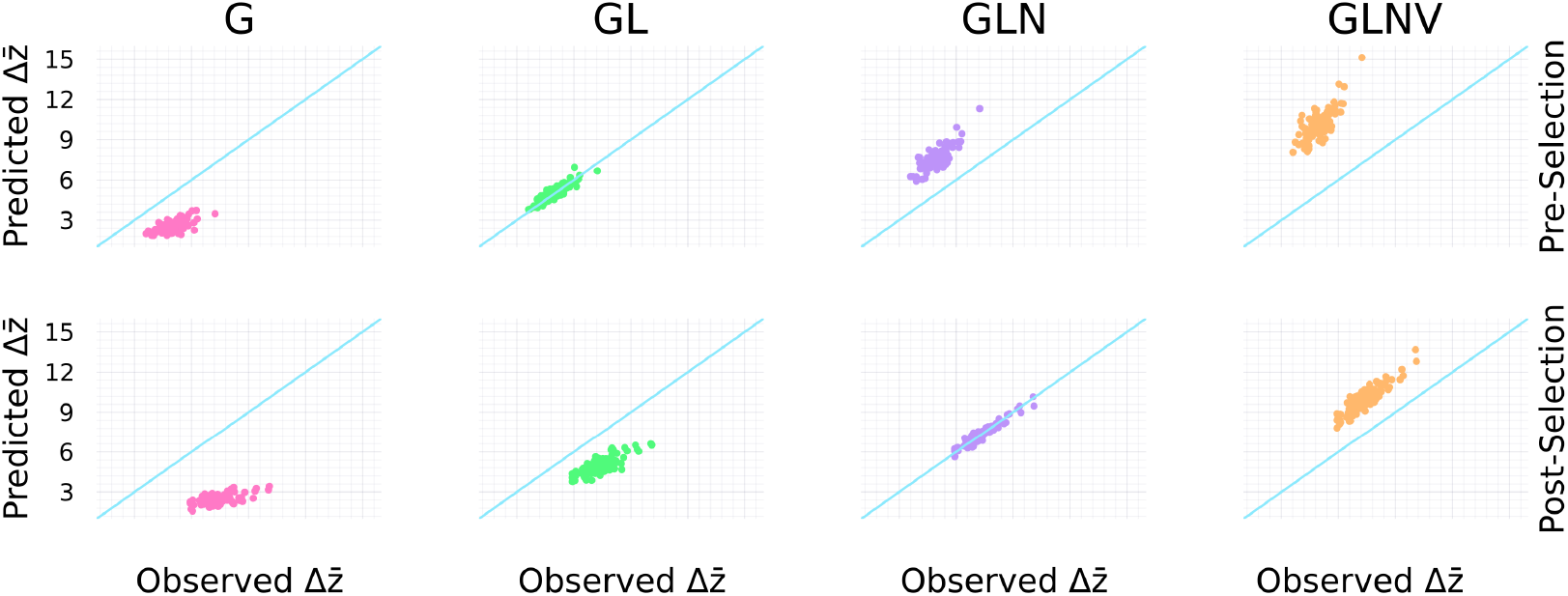
Comparison of change in host mean trait over a single consecutive generation 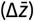 observed from simulations (x-axis) to that predicted by including different components of the host microbiome (y-axis). Each dot corresponds to an average over repeated runs for a given set of randomly drawn additive effects. Colors and column labels follow the same pairing described in the caption for Figure 3. Blue lines have unit slope and zero intercept. This figure demonstrates that 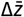 is *overestimated* when *including* either novel microbes (GLNV in top and bottom rows) or non-lineal microbes acquired from pre-selected parents (GLN in top row), is *underestimated* when *excluding* either all microbes (G in top and bottom rows) or non-lineal microbes acquired from post-selected parents (GL in bottom row). Predictions match observations on average when including host genes and lineal microbes, but excluding non-lineal microbes acquired from pre-selected parents and novel microbes (GL in top row), and when including only host genes, lineal microbes, and non-lineal microbes acquired from post-selected parents (GLN in bottom row).

#### 3.1.2 Simulation results

Our simulation results (summarized in Figure 5) demonstrate that when a host trait is significantly mediated by heritable microbes (such as lineal and possibly non-lineal microbes), then the classical breeders equation, which only accounts for genetic factors, severely underestimates the response to selection. In particular, for microbiome mediated traits, we find that the observed response to selection exceeds predictions based solely on host allele counts: 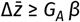.

Conversely, our model shows that including all trait-mediating microbes in the additive variance substantially overestimates the selection response. This result occurs because many microbes will be novel and thus not transmitted across host generations (which explains why the lines associated with GLNV and GLN coincide for both panels of Figure 3). More precisely, including trait mediating microbial taxa that are not transmitted across host generations (especially novel microbes, but possibly also non-lineal microbes) inflates the additive microbial variance which leads to observations that are exceeded by predictions: 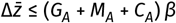.

In general, results from our simulations agree with our expectations described above. That is, by including *only* the transmissible microbes that contribute to an inter-generational response to selection on host individuals, we obtain the heuristic 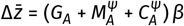.

The dichotomy of whether non-lineal microbes contribute to a selection response is artificial by design. More realistically, because different microbe individuals of the same taxon in a given host microbiome may have different patterns of ancestral associations with their hosts, microbial taxa cannot be neatly categorized by ancestral concordance as done here. Additionally, non-lineal microbe individuals may vary in the degree to which they contribute to a selection response because they may be transmitted either before or after the parental population experiences selection depending on the life-histories of the hosts and microbes. Such within-taxon variation could be captured in future work by modeling individual microbes’ ancestral concordance as draws from a taxon-specific distribution. Lastly, it is also possible that microbes can be transmitted beyond consecutive generations, but under our simulation model we expect the response to selection to be further attenuated in such cases.

### 3.2 Heritabilities and transmissibilities

Results from the previous sections provide insights into formulating a definition of microbial transmissibility. In particular, the indices we introduce here assume only microbes acquired from sources that contribute to a response to host-level selection (such as the lineal and possibly non-lineal microbes, see section 3.1) are included as trait mediating factors while calculating components of additive trait variation. Then, just as the *narrow-sense genetic heritability*, defined as *h*^2^ = *G*_*A*_/*P*, is useful for predicting the response to selection and can be measured using parent-offspring correlations, we define the *narrow-sense microbial transmissibility* as 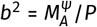, which is also useful for predicting the response to selection. However, unlike *h*^2^, only the lineal component of *b*^2^ will be measurable from parent-offspring correlations. If non-lineal microbes contribute to a response to selection, then their contribution to *b*^2^ may be quantified by first identifying the non-lineal donor-recipient pairs between the parent and offspring populations, and then measure correlations between donors and recipients. Of course, this approach to estimating *b*^2^ depends on our simplifying assumptions on the partitioning of host microbiome into lineal, non-lineal, and novel components. Because each taxa may consist of a mixture of lineal, non-lineal, and novel microbes, estimates of *b*^2^ will likely require a more sophisticated correlational analysis.

On their own, the narrow-sense heritability and narrow-sense microbial transmissibility are only useful for predicting a response to selection when the host trait is either entirely genetically mediated, or entirely microbially mediated. To obtain accurate predictions for the response to selection when host traits are partially genetically mediated and partially microbially mediated, we introduce notions of *total transmissibility*. These quantities are meant to capture all components of host trait variation explained by factors that facilitate a host response to selection (which we refer to as *selective factors* from hereon). Writing 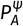 as the component of host trait variation explained only by additive effects of selective factors, we define the *narrow-sense total transmissibility* as 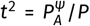. By focusing on additive effects, we can use the same general approach outlined in section 2.1 to quantify 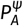. Here, because we assume the only selective factors are host genes and microbes, the narrow-sense total transmissibility becomes

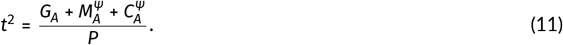

Hence, in general *t*^2^ ≠ *h*^2^ + *b*^2^. Instead, writing 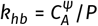, we can heuristically think of expanding the square (*h* + *b*)^2^ to arrive at *t*^2^ = *h*^2^ + *k*_*hb*_ + *b*^2^ where *k*_*hb*_ takes the place of the undefined symbol 2*hb*. Because *k*_*hb*_ is a covariance it can be positive or negative. So ignoring *k*_*hb*_ while calculating *t*^2^ can positively or negatively bias predictions for a microbiome mediated response to selection. Our work then suggests the form of the breeders equation introduced by Lush (1937), *R* = *h*^2^*S* (where 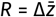 is the response to selection and *S* is the selection differential), naturally generalizes to *R* = *t*^2^*S* = (*h*^2^ + *k*_*hb*_ + *b*^2^)*S*. This reduces to the original expression in the absence of microbiome-mediated effects.

In our framework, broad-sense genetic heritability retains the same definition from classical quantitative genetics. In particular, the genotypic value *z*_*g*_ is the genotypic-microbic value *z*_*gm*_ averaged over host microbiomes. Then the *broad-sense genetic heritability* is defined as the proportion of host trait variance explained by genotypic variation: *H*^2^ = Var(*z*_*g*_)/*P*. In analogy, the *microbic value z*_*m*_ is the genotypic-microbic value *z*_*gm*_ averaged over host genotypes. To clarify that we are focusing specifically on host trait variation explained by microbes that contribute to a selection response, we propagate the superscript ^ψ^. In particular,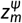 is *z*_*gm*_ averaged over host genotypes and components of host microbiomes that do not contain selective factors. This suggests that the *broad-sense microbial transmissibility* should be defined as the proportion of host trait variation explained by selective microbic variation: 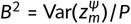. Setting *G*_*R*_ = Var(*z*) − *G* and 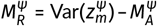 the residual variances left unexplained by additive genetic and selective additive microbial effects respectively, the broad-sense heritabilities can then be expressed as

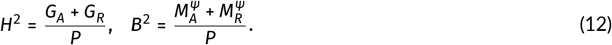

In general, the sum of these broad-sense heritabilities does not capture all of host trait variation explained by host genotype-microbiome pairs (*g, m*) because they do not account for covariances between allele counts and microbe abundances (quantified by 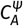) or for residual variation left unexplained by genotypic values and microbic values (quantified by 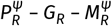, where 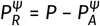). More generally, if the host trait is mediated by the selective factors *f* = (*f*_1_, …, *f*_*N*_), and *z*_*f*_ is the average trait value among hosts carrying factors *f*, then we define the *broad-sense total transmissibility* as the proportion of phenotypic variance explained by selective factors: *T* ^2^ = Var(*z*_*f*_)/*P*. Because we assume the only selective factors are host genes and microbes we have *z*_*f*_ = *z*_*gm*_, the genotypic-microbic value. Then, defining the symbol 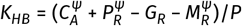, we obtain an expression for the broad-sense transmissibility as

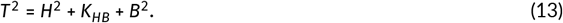

Again, we can intuitively think about calculating the expression for *T* ^2^ by expanding the square (*H* + *B*)^2^, with *K*_*HB*_ taking the place of the undefined symbol 2*HB*. Because 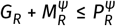, the component of *K*_*HB*_ due to differences in residual variation will always be non-negative. However, because 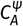 may be negative, *K*_*HB*_ will be negative when 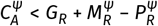. As a result, excluding *K*_*HB*_ from calculating *T*^2^ can positively or negatively bias estimates for the proportion of host trait variation explained by genetic and selective microbial factors.

The utility of the definition of narrow-sense total transmissibility introduced in this section for predicting a response to selection relies on a set of assumptions that simplify the process of microbial inheritance. Future work is needed to test their utility given the complexity of within host dynamics (Gerber, 2014), the microbiome assembly process (Costello et al., 2012), and host-host transmission (Sarkar et al., 2024).

## 4 DISCUSSION

There is growing evidence that microbes can be integral to the form, function, and fitness of larger organisms (Bordenstein and Theis, 2015). Microbes associated with plant and animal hosts have been shown to contribute to fundamental organismal processes, including development (McFall-Ngai et al., 2013), nutrition (David et al., 2013), pathogen protection (Koch and Schmid-Hempel, 2011) and even behavior (Cryan and Dinan, 2012). Microbes have been suggested to provide their plant and animal hosts with the capacity for rapid evolution (Rosenberg and Zilber-Rosenberg, 2016; Bisschop et al., 2022) and perhaps even to contribute to the evolutionary rescue of larger organisms from rapid environmental change (Pillai et al., 2016; Lennon et al., 2019).

This increasing interest in host-microbe interactions has led to a crucial need for a deeper understanding of how host-associated microbes influence the phenotypes of their hosts, and ultimately their evolution. But developing this understanding has been hampered by the lack of a comprehensive theoretical framework for considering the combined influences of host genetics, host-associated microbes and the surrounding environment on traits that emerge from such host-microbe systems, and corresponding statistical framework to disentangle these influences. Quantitative genetics is uniquely positioned to provide this foundation because it is focused on understanding the inheritance and evolution of the complex traits that result from multiple interacting sources of variation. While quantitative genetics was initially developed with genetic factors as the sole heritable units, it has since been extended to include factors subject to non-genetic inheritance, such as epigenetic alleles and cultural memes (Day and Bonduriansky, 2011; David and Ricard, 2019; Cavalli-Sforza and Feldman, 1981). Here we extend quantitative genetics by formally incorporating microbiomes as potentially heritable units that are similar to, but distinct from, host genetic factors.

Importantly, microbes are not merely their host’s *second genome*; there are fundamental differences between microbes in a microbiome and genes in a genome. These differences make applying quantitative genetic approaches directly to host microbiomes problematic. As a consequence, there is a need to expand theory of quantitative genetics to incorporate unique aspects of microbiome biology, as we have attempted to do here. We have built on previous efforts to establish quantitative genetic frameworks for microbiome-mediated traits (Henry et al., 2021) by generalizing the foundations of quantitative genetic theory. Our approach provides a formal definition for the component of phenotypic variation explained by additive effects of host genetic and microbial factors (*P*_*A*_), and how to partition this additive variance into additive genetic variance (*G*_*A*_) and the newly defined *additive microbial variance* (*M*_*A*_, the component of host trait variance explained by additive effects of microbe factors) and *additive gene-microbe covariance* (*C*_*A*_, which quantifies additive effects due to covariances between allele counts and microbe abundances across host individuals) such that *P*_*A*_ = *G*_*A*_ + *M*_*A*_ + *C*_*A*_.

Furthermore, to make accurate predictions for microbiome-mediated responses to selection on hosts, we found it necessary to introduce additional partitioning of host trait variation explained by host microbiomes. In particular, we suggest host microbiomes may be decomposed into three components: the *lineal* component which contains microbes passed along host lineages, the *novel* component which contains microbes acquired from the environment that have no previous associations with the host species, and the *non-lineal* component which contains all remaining microbes. We anticipate that whether or not microbes facilitate a response to selection depends on the microbiome component to which they belong. When microbial taxa that contribute to a selection response have been identified, we suggest quantifying 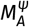 as the component of host trait variance explained *only* by the additive effects of those taxa. Using a simulation model, we find support for a generalized breeders equation taking the form 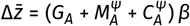. Building on this, we suggest definitions that generalize narrow-sense and broad-sense heritabilities to account for microbes that contribute to a selection response.

From this initial expansion, we can make a number of conclusions that are worth further theoretical and experimental exploration. For instance, Figures 3 and 4 together demonstrate that not all evolutionarily important microbes significantly correlate with host genes. This finding therefore negates a requirement of lineal transmission. In particular, Figure 3 shows that non-lineal microbes sourced from post-selected parents significantly contribute to a sustained response to selection over several host generations. In addition, Figure 4 shows that the within-host abundances of these same microbes do not significantly correlate with host allele counts (where significant here means to have greater magnitudes of correlations than novel microbe abundances). Another conclusion is in the converse direction; that not all microbes with abundances that significantly correlate with host allele counts contribute significantly to host trait variation. This result underscores the possibility that microbes transmitted across host generations do not necessarily mediate a response to host-level selection. For example, such correlations may occur when the host population is structured and exhibits random genetic drift. Furthermore, it is worth considering the degree to which processes analogous to classical random genetic drift emerge from microbiome-mediated host trait dynamics.

### Implications for association studies

Both of the conclusions in the preceding paragraph, that not all evolutionary important microbes correlate with host genes and not all microbes that correlate with host genes contribute to host trait variation, have important implications for how we study host-microbe interactions and their impact on host phenotype. The approaches commonly used for identifying the genes or microbes underlying a particular phenotype (e.g., GWAS or MWAS) require an understanding of how statistical associations form among phenotypes, genes and microbes (Camarinha-Silva et al., 2017; Difford et al., 2018; Qin et al., 2012; Pérez-Enciso et al., 2021). Because these associations are not necessarily causal, and because causal pathways may not be detected via statistical associations, there is a need for a more comprehensive theoretical foundation to guide approaches such as GWAS and MWAS. Importantly, microbiomes are dynamic ecological assemblages obeying different rules of inheritance than genomes, and so uncritical application of methods designed for GWAS may produce misleading results. The framework we introduce addresses this issue by clarifying when analogies between genes and microbes hold and when they break down, helping to replace false equivalences with a more accurate foundation for interpreting host–microbe trait associations. Furthermore, this framework offers a conceptual guide for interpreting MWAS results and for identifying microbial associations that are more likely to reflect biologically meaningful or evolutionarily relevant effects. Rather than assuming causality from variance explained, our framework points to ways of prioritizing microbial taxa based on patterns of shared ancestry with host lineages and association with host-level selection response, providing a clearer path for future empirical and statistical work.

In addition to extending the theoretical foundations of quantitative genetics, a key challenge moving forward will be to develop inference methods capable of estimating the host and microbial parameters defined in our framework. An interesting approach for future work will be to expand genomic regression approaches to include microbiomes for inferring parameters such as *T* ^2^ and *b*^2^ (Xavier et al., 2019). As with classical quantitative genetics, inference will require statistical models that account for relatedness among hosts and microbial lineages, and that distinguish causal from merely associated components. Importantly, as shown in recent work, failure to account for differences between causal loci and observed markers in genomic regression can lead to misleading heritability estimates (de los Campos et al., 2015). Analogous pitfalls are likely to occur in microbiome-based studies, underscoring the need for explicit statistical theory to accompany any expanded quantitative genetic framework.

### Implications for agriculture and wild populations

In livestock research, several studies have estimated the contribution of the gut microbiota to feed efficiency traits (Déru et al., 2022; Martinez-Boggio et al., 2024; Zhang et al., 2024) and in crops to stress tolerance and yield-related traits (Dwivedi et al., 2025; Mueller and Linksvayer, 2022), often by modeling microbial variation as an independent random effect alongside host genetic effects in linear mixed models. Unlike these empirical approaches, which frequently reduce dimensionality without biological interpretation (e.g., via principal components or OTU filtering thresholds), our framework provides biologically grounded decompositions based on patterns of host-microbe ancestral concordance. These decompositions may be estimated from lineage-traced or metagenomic data, and in doing so, provide a principled means of attributing trait variation to distinct classes of microbes. This allows our framework to complement and systematize empirical models used in breeding applications, while also offering a unified quantitative structure for comparing microbiome-mediated effects to other extended inheritance mechanisms. By explicitly modeling classes of microbial effects in parallel to classical host genetic effects, our approach can help disentangle microbiome contributions from confounded influences such as maternal effects, indirect genetic effects, and ecological inheritance, thereby clarifying their relative roles in shaping host phenotypes.

Beyond its relevance to agriculture, the framework we present here may also offer insights into how wild populations respond to natural selection. Long-term studies in wild systems have frequently documented cases where evolutionary change falls short of predictions based on observed heritability and directional selection (Kruuk et al., 2002; Pemberton, 2010; Pujol et al., 2018). Microbiome inheritance and its influence on host traits may contribute to this mismatch, but the mechanisms involved are likely complex and remain poorly understood. Different modes of microbial transmission and ecological dynamics could interact with host genetics in ways that violate standard quantitative genetic assumptions. Future work is needed to clarify how microbially mediated traits shape evolutionary predictions in natural populations. By formally integrating host-associated microbiomes into the foundations of quantitative genetic theory, our framework offers a potentially fruitful direction for addressing this challenge.

### Limitations and future directions

The framework presented throughout this paper has both general components that are independent of model assumptions, and specific components that are obtained by applying simplifying assumptions. In particular, both the approach to analysis of phenotypic variance in section 2.1 and our approach to partition host microbiomes by ancestral concordance in section 2.3.1 are independent of model assumptions. However, our results on the response of a microbiome mediated trait to selection (illustrated by Figure 5 and equations presented in the simulation results section) were obtained following a series of simplifying assumptions. These include independence of microbiome components (see section 2.3.2) and assumptions regarding the inheritance of microbiomes (summarized in Figure 2). In effect, our simulation model ignores within-host microbiome dynamics, the assembly process, and host-host microbe transmission, which has allowed us to focus on the effects of a novel inheritance mechanism on the dynamics of a microbiome-mediated quantitative character, and to identify the conditions under which distinct classes of microbes contribute to selection responses at the host level. Future work is needed to study models in which these assumptions are relaxed, and to apply our general framework of phenotypic variance analysis to both simulated and empirical data. More specifically, given the ecological nature of microbiomes, a fruitful avenue forward would be to interface our extension of quantitative genetics with metacommunity models of microbiome dynamics (Leibold et al., 2004; Koskella et al., 2017; Zeng et al., 2017; Miller et al., 2018). Such work can shed light on microbial ecological mechanisms that explain microbiome inheritance.

## 5 CONCLUSION

Microbes can be integral to the functioning of their animal and plant hosts, yet it is not well understood how microbiomes contribute to host fitness and evolution. This knowledge gap is due at least in part to the lack of a comprehensive theoretical framework for modeling how host-microbiome system-level traits emerge from the interactions of host genes, microbiome, and environment. We provide one such framework, by expanding theoretical quantitative genetics to include unique aspects of host-associated microbiomes, including multiple forms of Mendelian and non-Mendelian inheritance. This expansion leads to a formalization of several important concepts, including microbial heritabilities and transmissibilities, as well as fundamental quantities such as additive gene-microbe covariances. In addition, our framework provides an approach to partitioning quantitative trait variation into host-genetic and microbial components, allowing the theoretical exploration of how the joint contribution of host and microbiome to trait variation influences the evolution of host-microbiome systems. We consider our theoretical expansion as a first step toward a comprehensive incorporation of host-associated microbes into quantitative genetics.

## LITERATURE CITED

Ali, H. R. K., Hemeda, N. F. and Abdelaliem, Y. F. (2019) Symbiotic cellulolytic bacteria from the gut of the subterranean termite psammotermes hypostoma desneux and their role in cellulose digestion. AMB Express, 9. URL: 10.1186/s13568-019-0830-5.

Argaw-Denboba, A., Schmidt, T. S. B., Di Giacomo, M., Ranjan, B., Devendran, S., Mastrorilli, E., Lloyd, C. T., Pugliese, D., Paribeni, V., Dabin, J., Pisaniello, A., Espinola, S., Crevenna, A., Ghosh, S., Humphreys, N., Boruc, O., Sarkies, P., Zimmermann, M., Bork, P. and Hackett, J. A. (2024) Paternal microbiome perturbations impact offspring fitness. Nature, 629, 652–659. URL: 10.1038/s41586-024-07336-w.

Arora, J., Kinjo, Y., Šobotník, J., Buček, A., Clitheroe, C., Stiblik, P., Roisin, Y., Žifčáková, L., Park, Y. C., Kim, K. Y., Sillam-Dussès, D., Hervé, V., Lo, N., Tokuda, G., Brune, A. and Bourguignon, T. (2022) The functional evolution of termite gut microbiota. Microbiome, 10. URL: 10.1186/s40168-022-01258-3.

Awany, D. and Chimusa, E. R. (2020) Heritability jointly explained by host genotype and microbiome: will improve traits prediction? Briefings in Bioinformatics, 22. URL: 10.1093/bib/bbaa175.

Barton, N. and Turelli, M. (2004) Effects of genetic drift on variance components under a general model of epistasis. Evolution, 58, 2111–2132. URL: 10.1111/j.0014-3820.2004.tb01591.x.

Benson, A. K., Kelly, S. A., Legge, R., Ma, F., Low, S. J., Kim, J., Zhang, M., Oh, P. L., Nehrenberg, D., Hua, K., Kachman, S. D., Moriyama, E. N., Walter, J., Peterson, D. A. and Pomp, D. (2010) Individuality in gut microbiota composition is a complex polygenic trait shaped by multiple environmental and host genetic factors. Proceedings of the National Academy of Sciences, 107, 18933–18938. URL: 10.1073/pnas.1007028107.

Bijma, P., Muir, W. M. and Van Arendonk, J. A. M. (2007) Multilevel selection 1: Quantitative genetics of inheritance and response to selection. Genetics, 175, 277–288. URL: 10.1534/genetics.106.062711.

Bisschop, K., Kortenbosch, H. H., van Eldijk, T. J. B., Mallon, C. A., Salles, J. F., Bonte, D. and Etienne, R. S. (2022) Microbiome heritability and its role in adaptation of hosts to novel resources. Frontiers in Microbiology, 13. URL: 10.3389/fmicb.2022.703183.

Bonduriansky, R. and Day, T. (2009) Nongenetic inheritance and its evolutionary implications. Annual Review of Ecology, Evolution, and Systematics, 40, 103–125. URL: 10.1146/annurev.ecolsys.39.110707.173441.

Bordenstein, S. R. and Theis, K. R. (2015) Host biology in light of the microbiome: Ten principles of holobionts and hologenomes. PLOS Biology, 13, e1002226. URL: 10.1371/journal.pbio.1002226.

Bruijning, M., Henry, L. P., Forsberg, S. K. G., Metcalf, C. J. E. and Ayroles, J. F. (2021) Natural selection for imprecise vertical transmission in host–microbiota systems. Nature Ecology & Evolution, 6, 77–87. URL: 10.1038/s41559-021-01593-y.

Bürger, R. (2000) The mathematical theory of selection, recombination, and mutation. John Wiley & Sons.

Burns, A. R., Miller, E., Agarwal, M., Rolig, A. S., Milligan-Myhre, K., Seredick, S., Guillemin, K. and Bohannan, B. J. M. (2017) Interhost dispersal alters microbiome assembly and can overwhelm host innate immunity in an experimental zebrafish model. Proceedings of the National Academy of Sciences, 114, 11181–11186. URL: 10.1073/pnas.1702511114.

Camarinha-Silva, A., Maushammer, M., Wellmann, R., Vital, M., Preuss, S. and Bennewitz, J. (2017) Host genome influence on gut microbial composition and microbial prediction of complex traits in pigs. Genetics, 206, 1637–1644. URL: 10.1534/genetics.117.200782.

de los Campos, G., Sorensen, D. and Gianola, D. (2015) Genomic heritability: What is it? PLOS Genetics, 11, e1005048. URL: 10.1371/journal.pgen.1005048.

Cavalli-Sforza, L. L. and Feldman, M. W. (1981) Cultural transmission and evolution: A quantitative approach. No. 16. Princeton University Press.

Cheverud, J. M. and Routman, E. J. (1996) Epistasis as a source of increased additive genetic variance at population bottlenecks. Evolution, 50, 1042–1051. URL: 10.1111/j.1558-5646.1996.tb02345.x.

Costello, E. K., Stagaman, K., Dethlefsen, L., Bohannan, B. J. M. and Relman, D. A. (2012) The application of ecological theory toward an understanding of the human microbiome. Science, 336, 1255–1262. URL: 10.1126/science.1224203.

Cryan, J. F. and Dinan, T. G. (2012) Mind-altering microorganisms: the impact of the gut microbiota on brain and behaviour. Nature Reviews Neuroscience, 13, 701–712. URL: 10.1038/nrn3346.

Danchin, E., Charmantier, A., Champagne, F. A., Mesoudi, A., Pujol, B. and Blanchet, S. (2011) Beyond dna: integrating inclusive inheritance into an extended theory of evolution. Nature Reviews Genetics, 12, 475–486. URL: 10.1038/nrg3028.

David, I. and Ricard, A. (2019) A unified model for inclusive inheritance in livestock species. Genetics, 212, 1075–1099. URL: 10.1534/genetics.119.302375.

David, L. A., Maurice, C. F., Carmody, R. N., Gootenberg, D. B., Button, J. E., Wolfe, B. E., Ling, A. V., Devlin, A. S., Varma, Y., Fischbach, M. A., Biddinger, S. B., Dutton, R. J. and Turnbaugh, P. J. (2013) Diet rapidly and reproducibly alters the human gut microbiome. Nature, 505, 559–563. URL: 10.1038/nature12820.

Day, T. and Bonduriansky, R. (2011) A unified approach to the evolutionary consequences of genetic and nongenetic inheritance. The American Naturalist, 178, E18–E36. URL: 10.1086/660911.

Difford, G. F., Plichta, D. R., Løvendahl, P., Lassen, J., Noel, S. J., Højberg, O., Wright, A.-D. G., Zhu, Z., Kristensen, L., Nielsen, H. B., Guldbrandtsen, B. and Sahana, G. (2018) Host genetics and the rumen microbiome jointly associate with methane emissions in dairy cows. PLOS Genetics, 14, e1007580. URL: 10.1371/journal.pgen.1007580.

Dwivedi, S. L., Vetukuri, R. R., Kelbessa, B. G., Gepts, P., Heslop-Harrison, P., Araujo, A. S., Sharma, S. and Ortiz, R. (2025) Exploitation of rhizosphere microbiome biodiversity in plant breeding. Trends in Plant Science. URL: 10.1016/j.tplants.2025.04.004.

Déru, V., Tiezzi, F., Carillier-Jacquin, C., Blanchet, B., Cauquil, L., Zemb, O., Bouquet, A., Maltecca, C. and Gilbert, H. (2022) Gut microbiota and host genetics contribute to the phenotypic variation of digestive and feed efficiency traits in growing pigs fed a conventional and a high fiber diet. Genetics Selection Evolution, 54. URL: 10.1186/s12711-022-00742-6.

Fisher, R. A. (1918) Xv.—the correlation between relatives on the supposition of mendelian inheritance. Transactions of the Royal Society of Edinburgh, 52, 399–433. URL: 10.1017/S0080456800012163.

Fisher, R. A. (1941) Average excess and average effect of a gene substitution. Annals of Eugenics, 11, 53–63. URL: 10.1111/j.1469-1809.1941.tb02272.x.

Gerber, G. K. (2014) The dynamic microbiome. FEBS Letters, 588, 4131–4139. URL: 10.1016/j.febslet.2014.02.037.

Goodrich, J. K., Davenport, E. R., Waters, J. L., Clark, A. G. and Ley, R. E. (2016) Cross-species comparisons of host genetic associations with the microbiome. Science, 352, 532–535. URL: 10.1126/science.aad9379.

Gould, A. L., Zhang, V., Lamberti, L., Jones, E. W., Obadia, B., Korasidis, N., Gavryushkin, A., Carlson, J. M., Beerenwinkel, N. and Ludington, W. B. (2018) Microbiome interactions shape host fitness. Proceedings of the National Academy of Sciences, 115. URL: 10.1073/pnas.1809349115.

Helanterä, H. and Uller, T. (2010) The price equation and extended inheritance. Philosophy and Theory in Biology, 2. URL: 10.3998/ptb.6959004.0002.001.

Henry, L. P., Bruijning, M., Forsberg, S. K. G. and Ayroles, J. F. (2021) The microbiome extends host evolutionary potential. Nature Communications, 12. URL: 10.1038/s41467-021-25315-x.

Houwenhuyse, S., Stoks, R., Mukherjee, S. and Decaestecker, E. (2021) Locally adapted gut microbiomes mediate host stress tolerance. The ISME Journal, 15, 2401–2414. URL: 10.1038/s41396-021-00940-y.

Kirkpatrick, M. and Lande, R. (1989) The evolution of maternal characters. Evolution, 43, 485–503. URL: 10.1111/j.1558-5646.1989.tb04247.x.

Knights, D., Silverberg, M. S., Weersma, R. K., Gevers, D., Dijkstra, G., Huang, H., Tyler, A. D., van Sommeren, S., Imhann, F., Stempak, J. M., Huang, H., Vangay, P., Al-Ghalith, G. A., Russell, C., Sauk, J., Knight, J., Daly, M. J., Huttenhower, C. and Xavier, R. J. (2014) Complex host genetics influence the microbiome in inflammatory bowel disease. Genome Medicine, 6. URL: 10.1186/s13073-014-0107-1.

Koch, H. and Schmid-Hempel, P. (2011) Socially transmitted gut microbiota protect bumble bees against an intestinal parasite. Proceedings of the National Academy of Sciences, 108, 19288–19292. URL: 10.1073/pnas.1110474108.

Koskella, B., Hall, L. J. and Metcalf, C. J. E. (2017) The microbiome beyond the horizon of ecological and evolutionary theory. Nature Ecology & Evolution, 1, 1606–1615. URL: 10.1038/s41559-017-0340-2.

Kruuk, L. E. B., Slate, J., Pemberton, J. M., Brotherstone, S., Guinness, F. and Clutton-Brock, T. (2002) Antler size in red deer: Heritability and selection but no evolution. Evolution, 56, 1683–1695. URL: 10.1111/j.0014-3820.2002.tb01480.x.

Lande, R. (1976) Natural selection and random genetic drift in phenotypic evolution. Evolution, 314–334.

Leibold, M. A., Holyoak, M., Mouquet, N., Amarasekare, P., Chase, J. M., Hoopes, M. F., Holt, R. D., Shurin, J. B., Law, R., Tilman, D., Loreau, M. and Gonzalez, A. (2004) The metacommunity concept: a framework for multi-scale community ecology. Ecology Letters, 7, 601–613. URL: 10.1111/j.1461-0248.2004.00608.x.

Lennon, J. T., Mueller, E. A., Wisnoski, N. I. and Peralta, A. L. (2019) Microbial rescue effects: how microbiomes can save hosts from extinction. URL: 10.31219/osf.io/snr49.

Lush, J. (1937) Animal Breeding Plans. Collegiate Press, Incorporated.

Lynch, M. and Walsh, B. (1998) Genetics and analysis of quantitative traits, vol. 1. Sinauer Sunderland, MA.

Mackay, T. F. C., Stone, E. A. and Ayroles, J. F. (2009) The genetics of quantitative traits: challenges and prospects. Nature Reviews Genetics, 10, 565–577. URL: 10.1038/nrg2612.

Martinez-Boggio, G., Monteiro, H. F., Lima, F. S., Figueiredo, C. C., Bisinotto, R. S., Santos, J. E. P., Mion, B., Schenkel, F. S., Ribeiro, E. S., Weigel, K. A., Rosa, G. J. M. and Peñagaricano, F. (2024) Revealing host genome–microbiome networks underlying feed efficiency in dairy cows. Scientific Reports, 14. URL: 10.1038/s41598-024-77782-z.

Martiny, J. B. H., Jones, S. E., Lennon, J. T. and Martiny, A. C. (2015) Microbiomes in light of traits: A phylogenetic perspective. Science, 350. URL: 10.1126/science.aac9323.

Maurice, N. and Erdei, L. (2018) Termite Gut Microbiome, 69–99. Springer International Publishing. URL: 10.1007/978-3-319-72110-1_4.

McFall-Ngai, M. (2007) Care for the community. Nature, 445, 153–153. URL: 10.1038/445153a.

McFall-Ngai, M., Hadfield, M. G., Bosch, T. C. G., Carey, H. V., Domazet-Lošo, T., Douglas, A. E., Dubilier, N., Eberl, G., Fukami, T., Gilbert, S. F., Hentschel, U., King, N., Kjelleberg, S., Knoll, A. H., Kremer, N., Mazmanian, S. K., Metcalf, J. L., Nealson, K., Pierce, N. E., Rawls, J. F., Reid, A., Ruby, E. G., Rumpho, M., Sanders, J. G., Tautz, D. and Wernegreen, J. J. (2013) Animals in a bacterial world, a new imperative for the life sciences. Proceedings of the National Academy of Sciences, 110, 3229–3236. URL: 10.1073/pnas.1218525110.

Miller, E. T., Svanbäck, R. and Bohannan, B. J. (2018) Microbiomes as metacommunities: Understanding host-associated microbes through metacommunity ecology. Trends in Ecology & Evolution, 33, 926–935. URL: 10.1016/j.tree.2018.09.002.

Moore, A. J., Brodie, E. D. and Wolf, J. B. (1997) Interacting phenotypes and the evolutionary process: I. direct and indirect genetic effects of social interactions. Evolution, 51, 1352–1362. URL: 10.1111/j.1558-5646.1997.tb01458.x.

Morris, A. H. and Bohannan, B. J. M. (2024) Estimates of microbiome heritability across hosts. Nature Microbiology, 9, 3110–3119. URL: 10.1038/s41564-024-01865-w.

Mueller, U. G. and Linksvayer, T. A. (2022) Microbiome breeding: conceptual and practical issues. Trends in Microbiology, 30, 997–1011. URL: 10.1016/j.tim.2022.04.003.

Odling-Smee, J., Erwin, D. H., Palkovacs, E. P., Feldman, M. W. and Laland, K. N. (2013) Niche construction theory: A practical guide for ecologists. The Quarterly Review of Biology, 88, 3–28. URL: 10.1086/669266.

Opstal, E. J. v. and Bordenstein, S. R. (2015) Rethinking heritability of the microbiome. Science, 349, 1172–1173. URL: 10.1126/science.aab3958.

Org, E., Parks, B. W., Joo, J. W. J., Emert, B., Schwartzman, W., Kang, E. Y., Mehrabian, M., Pan, C., Knight, R., Gunsalus, R., Drake, T. A., Eskin, E. and Lusis, A. J. (2015) Genetic and environmental control of host-gut microbiota interactions. Genome Research, 25, 1558–1569. URL: 10.1101/gr.194118.115.

O’Brien, A. M., Laurich, J. R. and Frederickson, M. E. (2023) Evolutionary consequences of microbiomes for hosts: impacts on host fitness, traits, and heritability. Evolution, 78, 237–252. URL: 10.1093/evolut/qpad183.

Pemberton, J. M. (2010) Evolution of quantitative traits in the wild: mind the ecology. Philosophical Transactions of the Royal Society B: Biological Sciences, 365, 2431–2438. URL: 10.1098/rstb.2010.0108.

Petersen, C., Hamerich, I. K., Adair, K. L., Griem-Krey, H., Torres Oliva, M., Hoeppner, M. P., Bohannan, B. J. M. and Schulenburg, H. (2023) Host and microbiome jointly contribute to environmental adaptation. The ISME Journal, 17, 1953–1965. URL: 10.1038/s41396-023-01507-9.

Pillai, P., Gouhier, T. C. and Vollmer, S. V. (2016) Ecological rescue of host-microbial systems under environmental change. Theoretical Ecology, 10, 51–63. URL: 10.1007/s12080-016-0310-3.

Provine, W. B. (2001) The origins of theoretical population genetics: with a new afterword. University of Chicago Press.

Pujol, B., Blanchet, S., Charmantier, A., Danchin, E., Facon, B., Marrot, P., Roux, F., Scotti, I., Teplitsky, C., Thomson, C. E. and Winney, I. (2018) The missing response to selection in the wild. Trends in Ecology & Evolution, 33, 337–346. URL: 10.1016/j.tree.2018.02.007.

Pérez-Enciso, M., Zingaretti, L. M., Ramayo-Caldas, Y. and de los Campos, G. (2021) Opportunities and limits of combining microbiome and genome data for complex trait prediction. Genetics Selection Evolution, 53. URL: 10.1186/s12711-021-00658-7.

Qin, J., Li, Y., Cai, Z., Li, S., Zhu, J., Zhang, F., Liang, S., Zhang, W., Guan, Y., Shen, D., Peng, Y., Zhang, D., Jie, Z., Wu, W., Qin, Y., Xue, W., Li, J., Han, L., Lu, D., Wu, P., Dai, Y., Sun, X., Li, Z., Tang, A., Zhong, S., Li, X., Chen, W., Xu, R., Wang, M., Feng, Q., Gong, M., Yu, J., Zhang, Y., Zhang, M., Hansen, T., Sanchez, G., Raes, J., Falony, G., Okuda, S., Almeida, M., LeChatelier, E., Renault, P., Pons, N., Batto, J.-M., Zhang, Z., Chen, H., Yang, R., Zheng, W., Li, S., Yang, H., Wang, J., Ehrlich, S. D., Nielsen, R., Pedersen, O., Kristiansen, K. and Wang, J. (2012) A metagenome-wide association study of gut microbiota in type 2 diabetes. Nature, 490, 55–60. URL: 10.1038/nature11450.

Rice, S. (2004) Evolutionary Theory: Mathematical and Conceptual Foundations. Sinauer. URL: https://books.google.com/books?id=nGpoQgAACAAJ.

Rosenberg, E. and Zilber-Rosenberg, I. (2016) Microbes drive evolution of animals and plants: the hologenome concept. mBio, 7. URL: 10.1128/mBio.01395-15.

Rosenberg, E. and Zilber-Rosenberg, I. (2018) The hologenome concept of evolution after 10 years. Microbiome, 6. URL: 10.1186/s40168-018-0457-9.

Rosshart, S. P., Vassallo, B. G., Angeletti, D., Hutchinson, D. S., Morgan, A. P., Takeda, K., Hickman, H. D., McCulloch, J. A., Badger, J. H., Ajami, N. J., Trinchieri, G., Pardo-Manuel de Villena, F., Yewdell, J. W. and Rehermann, B. (2017) Wild mouse gut microbiota promotes host fitness and improves disease resistance. Cell, 171, 1015–1028.e13. URL: 10.1016/j.cell.2017.09.016.

Rothschild, D., Weissbrod, O., Barkan, E., Kurilshikov, A., Korem, T., Zeevi, D., Costea, P. I., Godneva, A., Kalka, I. N., Bar, N., Shilo, S., Lador, D., Vila, A. V., Zmora, N., Pevsner-Fischer, M., Israeli, D., Kosower, N., Malka, G., Wolf, B. C., Avnit-Sagi, T., Lotan-Pompan, M., Weinberger, A., Halpern, Z., Carmi, S., Fu, J., Wijmenga, C., Zhernakova, A., Elinav, E. and Segal, E. (2018) Environment dominates over host genetics in shaping human gut microbiota. Nature, 555, 210–215. URL: 10.1038/nature25973.

Roughgarden, J. (2023) Holobiont evolution: Population theory for the hologenome. The American Naturalist, 201, 763–778. URL: 10.1086/723782.

Roughgarden, J., Gilbert, S. F., Rosenberg, E., Zilber-Rosenberg, I. and Lloyd, E. A. (2017) Holobionts as units of selection and a model of their population dynamics and evolution. Biological Theory, 13, 44–65. URL: 10.1007/s13752-017-0287-1.

Russell, S. L. (2019) Transmission mode is associated with environment type and taxa across bacteria-eukaryote symbioses: a systematic review and meta-analysis. FEMS Microbiology Letters, 366. URL: 10.1093/femsle/fnz013.

Sandoval-Motta, S., Aldana, M. and Frank, A. (2017a) Evolving ecosystems: Inheritance and selection in the light of the microbiome. Archives of Medical Research, 48, 780–789. URL: 10.1016/j.arcmed.2018.01.002.

Sandoval-Motta, S., Aldana, M., Martínez-Romero, E. and Frank, A. (2017b) The human microbiome and the missing heritability problem. Frontiers in Genetics, 8. URL: 10.3389/fgene.2017.00080.

Sarkar, A., McInroy, C. J., Harty, S., Raulo, A., Ibata, N. G., Valles-Colomer, M., Johnson, K. V.-A., Brito, I. L., Henrich, J., Archie, E. A., Barreiro, L. B., Gazzaniga, F. S., Finlay, B. B., Koonin, E. V., Carmody, R. N. and Moeller, A. H. (2024) Microbial transmission in the social microbiome and host health and disease. Cell, 187, 17–43. URL: 10.1016/j.cell.2023.12.014.

Spor, A., Koren, O. and Ley, R. (2011) Unravelling the effects of the environment and host genotype on the gut microbiome. Nature Reviews Microbiology, 9, 279–290. URL: 10.1038/nrmicro2540.

Strang, G. et al. (2019) Linear algebra and learning from data, vol. 4. Wellesley-Cambridge Press Cambridge.

Tavalire, H. F., Christie, D. M., Leve, L. D., Ting, N., Cresko, W. A. and Bohannan, B. J. M. (2021) Shared environment and genetics shape the gut microbiome after infant adoption. mBio, 12. URL: 10.1128/mBio.00548-21.

Theis, K. R. (2018) Hologenomics: Systems-level host biology. mSystems, 3. URL: 10.1128/mSystems00164-17.

Tierney, B. T., Yang, Z., Luber, J. M., Beaudin, M., Wibowo, M. C., Baek, C., Mehlenbacher, E., Patel, C. J. and Kostic, A. D. (2019) The landscape of genetic content in the gut and oral human microbiome. Cell Host & Microbe, 26, 283–295.e8. URL: 10.1016/j.chom.2019.07.008.

Uller, T. and Helanterä, H. (2013) Non-genetic inheritance in evolutionary theory: a primer. Non-Genetic Inheritance, 1. URL: 10.2478/ngi-2013-0003.

Visscher, P. M., Hill, W. G. and Wray, N. R. (2008) Heritability in the genomics era — concepts and misconceptions. Nature Reviews Genetics, 9, 255–266. URL: 10.1038/nrg2322.

van Vliet, S. and Doebeli, M. (2019) The role of multilevel selection in host microbiome evolution. Proceedings of the National Academy of Sciences, 116, 20591–20597. URL: 10.1073/pnas.1909790116.

Walsh, B. and Lynch, M. (2018) Evolution and selection of quantitative traits. Oxford University Press.

Wang, J., Chen, L., Zhao, N., Xu, X., Xu, Y. and Zhu, B. (2018) Of genes and microbes: solving the intricacies in host genomes. Protein & Cell, 9, 446–461. URL: 10.1007/s13238-018-0532-9.

Week, B., Russel, S., Schulenburg, H., Bohannan, B. and Bruijning, M. (2024) Understanding host-microbiome evolution through the lens of evolutionary theory: New tricks for old dogs. URL: 10.32942/X2H055.

Wendling, C. C. and Wegner, K. M. (2015) Adaptation to enemy shifts: rapid resistance evolution to local vibrio spp. in invasive pacific oysters. Proceedings of the Royal Society B: Biological Sciences, 282, 20142244. URL: 10.1098/rspb.2014.2244.

Wilson, A. J., Réale, D., Clements, M. N., Morrissey, M. M., Postma, E., Walling, C. A., Kruuk, L. E. B. and Nussey, D. H. (2009) An ecologist’s guide to the animal model. Journal of Animal Ecology, 79, 13–26. URL: 10.1111/j.1365-2656.2009.01639.x.

Xavier, A., Muir, W. M. and Rainey, K. M. (2019) bwgr: Bayesian whole-genome regression. Bioinformatics, 36, 1957–1959. URL: 10.1093/bioinformatics/btz794.

Zeng, Q., Wu, S., Sukumaran, J. and Rodrigo, A. (2017) Models of microbiome evolution incorporating host and microbial selection. Microbiome, 5. URL: 10.1186/s40168-017-0343-x.

Zhang, W., Lan, F., Zhou, Q., Gu, S., Li, X., Wen, C., Yang, N. and Sun, C. (2024) Host genetics and gut microbiota synergistically regulate feed utilization in egg-type chickens. Journal of Animal Science and Biotechnology, 15. URL: 10.1186/s40104-024-01076-7.

Zilber-Rosenberg, I. and Rosenberg, E. (2008) Role of microorganisms in the evolution of animals and plants: the hologenome theory of evolution. FEMS Microbiology Reviews, 32, 723–735. URL: 10.1111/j.1574-6976.2008.00123.x.

