## Supplementary Material for "Quantitative Genetics of Microbiome Mediated Traits"

CONTENTS

1 Justification of Variance Partitioning 1

2 Additive Variances under Pairwise Interactions 2

3 Tables of Parameter Values used in Simulations 5

### 1 | JUSTIFICATION OF VARIANCE PARTITIONING

Here we provide calculations to justify the partitioning of variance introduced in section 2.1 of the main text. The calculations here hold for any collection of numerically coded factors $f = (f_1, \dots, f_n)^\dagger$. In the main text, the $f_i$ are equal to the allele counts $g_i$ for $i = 1, \dots, L$, and are equal to relative microbial abundances $m_i$ for $i = L+1, \dots, L+S$. Instead of the additive effect specific to genotype-microbiome pairs $\alpha_{gm}$, we focus on obtaining an expression for the additive effect for the vector of arbitrary factors $f$, denoted $\alpha_f$. The additivity constraint on $\alpha_f$ implies $\alpha_f = \alpha_0 + \sum_i \alpha_i f_i$, which we abbreviate to $\alpha_f = \alpha_0 + \langle \alpha, f \rangle$ with $\alpha = (\alpha_1, \dots, \alpha_n)^\dagger$ the vector of additive effects associated with each component of $f$, and $\alpha_0$ is the intercept of the linear model. In the main text $\alpha_i = \gamma_i$ for $i = 1, \dots, L$ and $\alpha_i = \omega_i$ for $i = L+1, \dots, L+S$. We make the decomposition $z_f - \bar{z} = \delta_f = \alpha_f + \rho_f$, i.e., $\rho$ is the residual. Following the definition of $P_R$ in the main text we have $P_R = \sum_f p_f \rho_f^2$, where $p_f$ is the frequency of $f$ in the population. Then $P_R$ can be rewritten as $P_R = \sum_f p_f (\delta_f - \alpha_f)^2$. Our goal is to minimize $P_R$. Partial differentiation with respect to the intercept returns

$$\partial_{\alpha_0} P_R = -2 \sum_f p_f (\delta_f - \alpha_0 - \langle \alpha, f \rangle) = -2 \sum_f p_f \rho_f = -2\bar{\rho}. \quad (1)$$

The optimality condition $\partial_{\alpha_0} P_R = 0$ then implies $\bar{\rho} = 0$, and hence we can set $\alpha_0 = -\sum_f p_f \langle \alpha, f \rangle = -\langle \alpha, \bar{f} \rangle = -\sum_i \alpha_i \bar{f}_i$. Partial differentiation with respect to additive effect $\alpha_i$ for $i = 1, \dots, n$ returns

$$\partial_{\alpha_i} P_R = -2 \sum_f p_f (\delta_f - \alpha_0 - \langle \alpha, f \rangle) f_i = -2 \sum_f p_f \rho_f f_i = -2\,\mathrm{Cov}(\rho, f_i), \quad (2)$$

where the last equality holds because $\bar{\rho} = 0$. Then $\partial_{\alpha_i} P_R = 0$ implies $\mathrm{Cov}(\rho, f_i) = 0$. The expression for $\partial_{\alpha_i} P_R$ can also be written as

$$\partial_{\alpha_i} P_R = -2\,\mathrm{Cov}(\delta_f, f_i) + 2\,\mathrm{Cov}(\langle \alpha, f \rangle, f_i), \quad (3)$$

Note that $\mathrm{Cov}(\delta_f, f_i) = \mathrm{Cov}(z_f, f_i)$ and $\mathrm{Cov}(\langle \alpha, f \rangle, f_i) = \sum_j \alpha_j \mathrm{Cov}(f_j, f_i)$. Hence, $\partial_{\alpha_i} P_R = 0$ also implies

$$\mathrm{Cov}(z_f, f_i) = \sum_j \alpha_j \mathrm{Cov}(f_i, f_j). \quad (4)$$

Writing $\mathrm{Cov}(z_f, f)$ as the vector with $i$th entry $\mathrm{Cov}(z_f, f_i)$ and $\Sigma$ as the $n \times n$ matrix with $i,j$th entry $\mathrm{Cov}(f_i, f_j)$, the expression above can be written in vector-matrix notation as $\mathrm{Cov}(z_f, f) = \Sigma\,\alpha$. Hence, $\alpha = \Sigma^{-1}\,\mathrm{Cov}(z_f, f)$ when $\Sigma^{-1}$ exists and $\alpha_0 = -\langle \alpha, \bar{f} \rangle$ with $\bar{f} = (\bar{f}_1, \dots, \bar{f}_n)^\dagger$. This establishes the definition of additive genetic and microbial effects provided in the main text when $\Sigma$ is non-singular.

The result $\mathrm{Cov}(\rho_f, f_i) = 0$ implies $\sum_f p_f \alpha_f \rho_f = 0$, which then formally establishes the partitioning of the variance of the marginal value $\mathrm{Var}(z_f) = \sum_f p_f (z_f - \bar{z})^2$ as $\mathrm{Var}(z_f) = P_A + P_R$, with $P_A = \sum_f p_f \alpha_f^2$. More generally, given $K$ different classes of factors such that $f = (f_1^1, \dots, f_{n_1}^1, \dots, f_1^K, \dots, f_{n_K}^K)^\dagger$ and $\alpha = (\alpha_1^1, \dots, \alpha_{n_1}^1, \dots, \alpha_1^K, \dots, \alpha_{n_K}^K)^\dagger$, the expression for $P_A$ can be rewritten as

$$P_A = \sum_{k=1}^{K} P_{A,k} + \sum_{k=1}^{K} \sum_{\ell=1}^{k-1} C_{A,k,\ell}, \quad (5)$$

where $P_{A,k} = \sum_f p_f \alpha_{f^k}^2$ and $C_{A,k,\ell} = 2\sum_f p_f \alpha_{f^k}\alpha_{f^\ell}$, with $f^k = (f_1^k, \dots, f_{n_k}^k)^\dagger$, $\alpha^k = (\alpha_1^k, \dots, \alpha_{n_k}^k)^\dagger$, and $\alpha_{f_k} = \alpha_0^k + \langle \alpha^k, f^k \rangle$. Here we have introduced new partition-specific intercepts, which must satisfy $\alpha_0 = \sum_k \alpha_0^k$, and so we can take $\alpha_0^k = -\langle \alpha^k, \bar{f}^k \rangle$. With this choice the contributions to $\alpha_f$ from each partition are centered, so the resulting partition of variance comes out in terms of variances and covariances (without this choice of $\alpha_0^k$ we would have sums of squares instead of (co)variances). This returns our partitionings introduced in section 2.1 and in section 2.3.2 as special cases. Finally, our choice for $\alpha_0^k$ implies

$$P_{A,k} = \sum_{i=1}^{n_K} (\alpha_i^k)^2 \operatorname{Var}(f_i^k) + 2 \sum_{i=1}^{n_k} \sum_{j=1}^{i-1} \alpha_i^k \alpha_j^k \operatorname{Cov}(f_i^k, f_j^k), \tag{6a}$$

$$C_{A,k,\ell} = 2 \sum_{i=1}^{n_k} \sum_{j=1}^{n_\ell} \alpha_i^k \alpha_j^\ell \operatorname{Cov}(f_i^k, f_j^\ell), \tag{6b}$$

and this returns equations (5) of the main text as a special case.

The above partitioning of $P_A$ is unique only when $\Sigma$ is non-singular. When $\Sigma$ is singular, the Moore–Penrose pseudoinverse of $\Sigma$ is commonly used in place of $\Sigma^{-1}$. This approach is useful for approximating $P_A$, but the estimate of $\alpha$ loses biological significance as a consequence of its lack of uniqueness.

### 2 | ADDITIVE VARIANCES UNDER PAIRWISE INTERACTIONS

Here we compute expressions for host trait variation using a model that assumes genotypic-microbic values are determined by mechanistic additive effects of host allele counts $g_1, \dots, g_L$ and microbial abundances $m_1, \dots, m_S$ in addition to mechanistic effects explained by products of host allele counts $g_i g_j$, products of microbial abundances $m_i m_j$, and products of host allele counts and microbial abundances $g_i m_j$. This model incorporates all possible pairwise products of trait mediating factors analyzed, and therefore generalizes the model presented in section 5 of the main text which focused only on products of host allele counts and microbial abundances. The model here can be expressed as

$$z_{gm} = z_0 + \sum_{i=1}^{L} \mathring{\gamma}_i g_i + \sum_{i=1}^{L} \sum_{j=1}^{i} \mathring{\gamma}_{ij} g_i g_j + \sum_{i=1}^{S} \mathring{\omega}_i m_i + \sum_{i=1}^{S} \sum_{j=1}^{i} \mathring{\omega}_{ij} m_i m_j + \sum_{i=1}^{L} \sum_{j=1}^{S} \mathring{\chi}_{ij} g_i m_j. \tag{7}$$

Our goal here is to compute $G_A = \sum_{gm} p_{gm} \gamma_g^2$, $M_A = \sum_{gm} p_{gm} \omega_g^2$, and $C_A = \sum_{gm} p_{gm} \gamma_g \omega_m$ given the above model under the assumption that all factors $g_1, \dots, g_L, m_1, \dots, m_S$ are linearly independent so that $\Sigma^{-1}$ exists (which implies uniqueness of $G_A, M_A, C_A$). To begin, note that

$$\begin{aligned} \operatorname{Cov}(z_{gm}, g_k) = &\sum_{i=1}^{L} \mathring{\gamma}_i \operatorname{Cov}(g_i, g_k) + \sum_{i=1}^{S} \mathring{\omega}_i \operatorname{Cov}(m_i, g_k) \\ &+ \sum_{i=1}^{L} \sum_{j=1}^{i} \mathring{\gamma}_{ij} \operatorname{Cov}(g_i g_j, g_k) + \sum_{i=1}^{S} \sum_{j=1}^{S} \mathring{\omega}_{ij} \operatorname{Cov}(m_i m_j, g_k) + \sum_{i=1}^{L} \sum_{j=1}^{S} \mathring{\chi}_{ij} \operatorname{Cov}(g_i m_j, g_k), \end{aligned} \tag{8a}$$

$$\begin{aligned}\mathrm{Cov}(z_{gm}, m_k) = \sum_{i=1}^{L} \mathring{\gamma}_i \mathrm{Cov}(g_i, m_k) + \sum_{i=1}^{S} \mathring{\omega}_i \mathrm{Cov}(m_i, m_k) \\ + \sum_{i=1}^{L}\sum_{j=1}^{i} \mathring{\gamma}_{ij} \mathrm{Cov}(g_i g_j, m_k) + \sum_{i=1}^{S}\sum_{j=1}^{i} \mathring{\omega}_{ij} \mathrm{Cov}(m_i m_j, m_k) + \sum_{i=1}^{L}\sum_{j=1}^{S} \mathring{\chi}_{ij} \mathrm{Cov}(g_i m_j, m_k). \qquad (8b)\end{aligned}$$

Writing $\mathring{\alpha} = (\mathring{\gamma}_1, \dots, \mathring{\gamma}_L, \mathring{\omega}_1, \dots, \mathring{\omega}_S)$, and $v$ the vector-valued function of $g$ and $m$ with $k$th entry

$$v_k = \sum_{i=1}^{L}\sum_{j=1}^{i} \mathring{\gamma}_{ij} \mathrm{Cov}(g_i g_j, g_k) + \sum_{i=1}^{S}\sum_{j=1}^{i} \mathring{\omega}_{ij} \mathrm{Cov}(m_i m_j, g_k) + \sum_{i=1}^{L}\sum_{j=1}^{S} \mathring{\chi}_{ij} \mathrm{Cov}(g_i m_j, g_k), \; k = 1, \dots, L, \qquad (9a)$$

$$v_k = \sum_{i=1}^{L}\sum_{j=1}^{i} \mathring{\gamma}_{ij} \mathrm{Cov}(g_i g_j, m_{k-L}) + \sum_{i=1}^{S}\sum_{j=1}^{i} \mathring{\omega}_{ij} \mathrm{Cov}(m_i m_j, m_{k-L}) + \sum_{i=1}^{L}\sum_{j=1}^{S} \mathring{\chi}_{ij} \mathrm{Cov}(g_i m_j, m_{k-L}), \; k = L+1, \dots, L+S, \qquad (9b)$$

we have $\mathrm{Cov}(z_{gm}, a) = \Sigma\mathring{\alpha} + v$. Hence $\alpha = \mathring{\alpha} + \Sigma^{-1} v$, and writing $u = \Sigma^{-1} v$ the vector-valued function of $g$ and $m$ resulting from a transformation on $v$, we have $\gamma_i = \mathring{\gamma}_i + u_i$ and $\omega_i = \mathring{\omega}_i + u_{i+L}$.

$$\gamma_g = \sum_{i=1}^{L} (\mathring{\gamma}_i + u_i) g_i, \quad \omega_g = \sum_{i=1}^{S} (\mathring{\omega}_i + u_{i+L}) m_i. \qquad (10a)$$

Consequently

$$\bar{\gamma} = \sum_{i=1}^{L} \mathring{\gamma}_i \bar{g}_i + \overline{g_i u_i}, \quad \bar{\omega} = \sum_{i=1}^{S} \mathring{\omega}_i \bar{m}_i + \overline{m_i u_{i+L}}, \qquad (11)$$

where $\overline{g_i u_i}$ and $\overline{m_i u_{i+L}}$ are averages of $g_i u_i$ and $m_i u_{i+L}$ across genotype-microbiome pairs in the host population. To simplify expressions we write $[x] = x - \bar{x}$ and $[xy] = xy - \overline{xy}$. This leads to

$$[\gamma_g]^2 = \sum_{i=1}^{L} \mathring{\gamma}_i^2 [g_i]^2 + 2\mathring{\gamma}_i [g_i][g_i u_i] + [g_i u_i]^2 + 2\sum_{i=1}^{L}\sum_{j=1}^{i-1} \mathring{\gamma}_i \mathring{\gamma}_j [g_i][g_j] + \mathring{\gamma}_i [g_i][g_j u_j] + \mathring{\gamma}_j [g_j][g_i u_i] + [g_i u_i][g_j u_j], \qquad (12a)$$

$$[\omega_m]^2 = \sum_{i=1}^{S} \mathring{\omega}_i^2 [m_i]^2 + 2\mathring{\omega}_i [m_i][m_i u_{i+L}] + [m_i u_{i+L}]^2 + 2\sum_{i=1}^{S}\sum_{j=1}^{i-1} \mathring{\omega}_i \mathring{\omega}_j [m_i][m_j] + \mathring{\omega}_i [m_i][m_j u_{j+L}] + \mathring{\omega}_j [m_j][m_i u_{i+L}] + [m_i u_{i+L}][m_j u_{j+L}] \qquad (12b)$$

$$[\gamma_g][\omega_m] = \sum_{i=1}^{L}\sum_{j=1}^{S} \mathring{\gamma}_i \mathring{\omega}_j [g_i][m_i] + \mathring{\gamma}_i [g_i][m_j u_{j+L}] + \mathring{\omega}_j [m_j][g_i u_i] + [g_i u_i][m_j u_{j+L}]. \qquad (12c)$$

We can therefore write

$$\begin{aligned}G_A = \sum_{i=1}^{L} \mathring{\gamma}_i^2 \,\mathrm{Var}(g_i) + 2\mathring{\gamma}_i \,\mathrm{Cov}(g_i, g_i u_i) + \mathrm{Var}(g_i u_i) \\ + 2\sum_{i=1}^{L}\sum_{j=1}^{i-1} \mathring{\gamma}_i \mathring{\gamma}_j \,\mathrm{Cov}(g_i, g_j) + \mathring{\gamma}_i \,\mathrm{Cov}(g_i, g_j u_j) + \mathring{\gamma}_j \,\mathrm{Cov}(g_j, g_i u_i) + \mathrm{Cov}(g_i u_i, g_j u_j), \qquad (13a)\end{aligned}$$

$$
\begin{aligned}
M_A = & \sum_{i=1}^{S} \mathring{\omega}_i^2 \operatorname{Var}(m_i) + 2\mathring{\omega}_i \operatorname{Cov}(m_i, m_i u_{i+L}) + \operatorname{Var}(m_i u_{i+L}) \\
& + 2 \sum_{i=1}^{S} \sum_{j=1}^{i-1} \mathring{\omega}_i \mathring{\omega}_j \operatorname{Cov}(m_i, m_j) + \mathring{\omega}_i \operatorname{Cov}(m_i, m_j u_{j+L}) + \mathring{\omega}_j \operatorname{Cov}(m_j, m_i u_{i+L}) + \operatorname{Cov}(m_i u_{i+L}, m_j u_{j+L}), \quad (13b)
\end{aligned}
$$

$$
C_A = 2 \sum_{i=1}^{L} \sum_{j=1}^{S} \mathring{\gamma}_i \mathring{\omega}_j \operatorname{Cov}(g_i, m_j) + \mathring{\gamma}_i \operatorname{Cov}(g_i, m_j u_{j+L}) + \mathring{\omega}_j \operatorname{Cov}(m_j, g_i u_i) + \operatorname{Cov}(g_i u_i, m_j u_{j+L}). \quad (13c)
$$

### 3 | TABLES OF PARAMETER VALUES USED IN SIMULATIONS

**TABLE 1** Parameter values used for simulating data presented in Figure 5 of main text

| ine Symbol | Meaning | Value |
|---|---|---|
| ine nr | Number of times to initialize parental population | 100 |
| sr | Number of times to repeat selection of hosts | 20 |
| or | Number of times to repeat production of offspring | 20 |
| $h^2$ | Expected proportion of additive genetic variance | 0.25 |
| $\lambda^2$ | Expected proportion of additive lineal variance | 0.25 |
| $\nu^2$ | Expected proportion of additive non-lineal variance | 0.25 |
| $\varepsilon^2$ | Expected proportion of additive external variance | 0.25 |
| L | Number of host genetic loci | 100 |
| SL | Number of lineal microbe taxa | 100 |
| SN | Number of non-lineal microbe taxa | 100 |
| SE | Number of external microbe taxa | 100 |
| $\mathbb{E}P$ | Expected total phenotypic variance | 100 |
| p | Probability of alternate allele at each locus in parental genomes | 0.5 |
| n | Size of host populations | 1000 |
| s | Selection coefficient | 0.001 |
| KL | Expected abundance for lineal microbe taxa | 50 |
| KN | Expected abundance for non-lineal microbe taxa | 50 |
| KE | Expected abundance for external microbe taxa | 50 |
| ine | | |

**TABLE 2** Parameter values used for simulating data presented in Figures 3 and 4 of main text

| ine Symbol | Meaning | Value |
|---|---|---|
| ine T | Number of host generations | 10 |
| ts_rep | Number of time series to repeat | 20 |
| nr | Number of times to initialize parental population | 50 |
| sr | Number of times to repeat selection of hosts | 10 |
| or | Number of times to repeat production of offspring | 10 |
| $h^2_{GLNE}$ | Expected proportion of additive genetic variance for GLNE | 0.25 |
| $\lambda^2_{GLNE}$ | Expected proportion of additive lineal variance for GLNE | 0.25 |
| $\nu^2_{GLNE}$ | Expected proportion of additive non-lineal variance for GLNE | 0.25 |
| $\varepsilon^2 GLNE$ | Expected proportion of additive external variance for GLNE | 0.25 |
| $h^2_{GLN}$ | Expected proportion of additive genetic variance for GLN | 0.33 |
| $\lambda^2_{GLN}$ | Expected proportion of additive lineal variance for GLN | 0.33 |
| $\nu^2_{GLN}$ | Expected proportion of additive non-lineal variance for GLN | 0.33 |
| $\varepsilon^2_{GLN}$ | Expected proportion of additive external variance for GLN | 1e-3 |
| $h^2_{GL}$ | Expected proportion of additive genetic variance for GL | 0.5 |
| $\lambda^2_{GL}$ | Expected proportion of additive lineal variance for GL | 0.5 |
| $\nu^2_{GL}$ | Expected proportion of additive non-lineal variance for GL | 1e-3 |
| $\varepsilon^2_{GL}$ | Expected proportion of additive external variance for GL | 1e-3 |
| $h^2_{G}$ | Expected proportion of additive genetic variance for G | 0.99 |
| $\lambda^2_{G}$ | Expected proportion of additive lineal variance for G | 1e-3 |
| $\nu^2_{G}$ | Expected proportion of additive non-lineal variance for G | 1e-3 |
| $\varepsilon^2_{G}$ | Expected proportion of additive external variance for G | 1e-3 |
| L | Number of host genetic loci | 100 |
| SL | Number of lineal microbe taxa | 100 |
| SN | Number of non-lineal microbe taxa | 100 |
| SE | Number of external microbe taxa | 100 |
| $\mathbb{E}[P_{GLNE}]$ | Expected total phenotypic variance for GLNE | 100 |
| $\mathbb{E}[P_{GLN}]$ | Expected total phenotypic variance for GLN | 75 |
| $\mathbb{E}[P_{GL}]$ | Expected total phenotypic variance for GL | 50 |
| $\mathbb{E}[P_{G}]$ | Expected total phenotypic variance for G | 25 |
| $p$ | Probability of alternate allele at each locus in parental genomes | 0.5 |
| $n$ | Size of host populations | 1000 |
| $s$ | Selection coefficient | 0.001 |
| KL | Expected abundance for lineal microbe taxa | 50 |
| KN | Expected abundance for non-lineal microbe taxa | 50 |
| KE | Expected abundance for external microbe taxa | 50 |
| ine | | |
